# The RecBCD complex interacts directly with the DNA sliding clamp in *Escherichia coli*

**DOI:** 10.1101/2025.08.20.671105

**Authors:** Ida Mathilde Marstein Riisnæs, Synnøve Brandt Ræder, Signe Simonsen, Krister Vikedal, Paul Hoff Backe, Line Johnsen, Magnar Bjørås, James Alexander Booth, Birthe B. Kragelund, Kirsten Skarstad, Emily Helgesen

**Author notes:** Co-first authors (Ida Mathilde Marstein Riisnæs, Synnøve Brandt Ræder). Current affiliation: Aker BioMarine, Human Health Ingredients AS, 1327 Lysaker, Norway.

## Abstract

DNA sliding clamps are essential coordinators of genome replication and maintenance across all domains of life and serve as platforms for recruiting diverse binding partners. The bacterial DNA sliding clamp, β-clamp, functions analogous to the eukaryotic proliferating cell nuclear antigen (PCNA), yet its full interactome remains incompletely characterized. Here, we identify and characterize a previously unrecognized interaction between β-clamp and the DNA repair helicase-nuclease complex RecBCD, specifically through its RecB subunit. Using bacterial two-hybrid assays, co-immunoprecipitation, and fluorescence microscopy, we show that RecB physically associates with β-clamp. Nuclear magnetic resonance (NMR) spectroscopy demonstrates binding to the canonical ligand binding pocket in β-clamp and identifies a clamp-binding motif within the RecB nuclease domain which targeted mutagenesis abolishes. Moreover, disrupting the interaction between RecB and β-clamp compromises *Escherichia coli* survival following DNA damage and impairs the DNA degradation activity of RecBCD. These findings expand the known β-clamp interactome and uncover a possible role for β-clamp in DNA double-strand break repair, offering new insights into bacterial DNA metabolism and potential avenues for therapeutic intervention.

**GRAPHICAL ABSTRACT:** 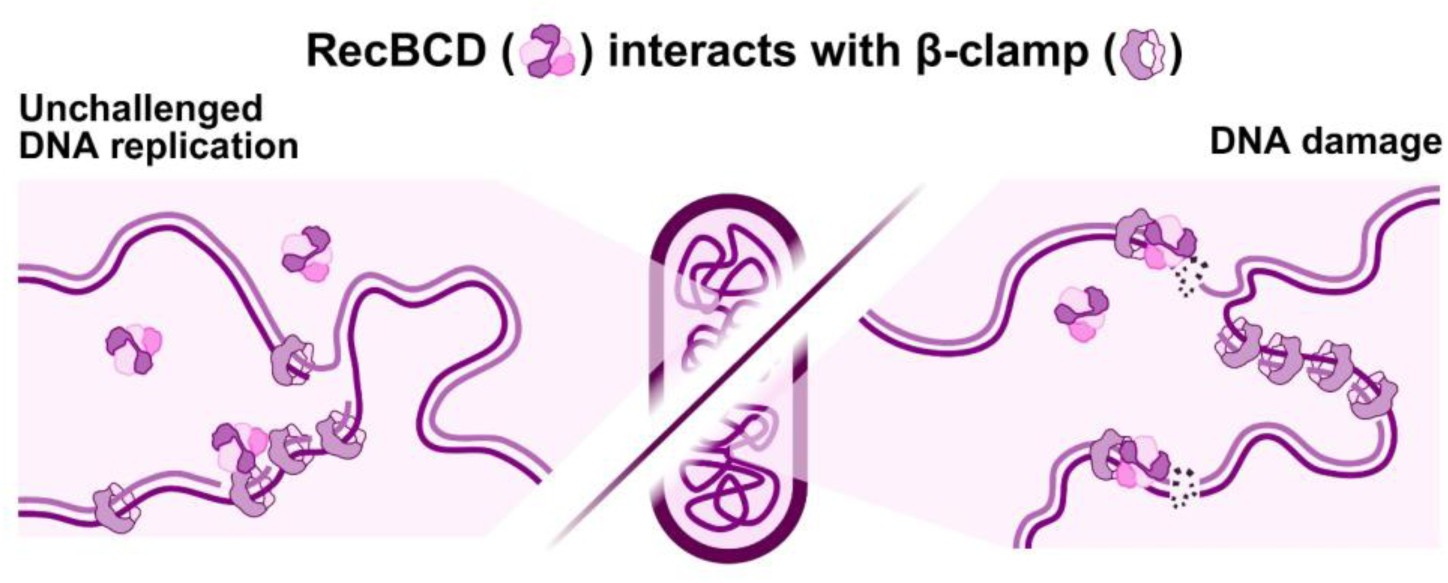

## INTRODUCTION

DNA sliding clamps are ring-shaped protein complexes fundamental to DNA replication and genome maintenance across all domains of life. In bacteria, the protein β-clamp (*dnaN*) serves as an essential hub protein, coordinating multiple cellular processes through interactions with diverse binding partners (1). As a processivity factor, it ensures efficient DNA replication by tethering DNA polymerases (Pol I, and the α and ε subunits of Pol III) to the DNA (2,3). β-clamp also engages with the replication initiation factor DnaA (4), its regulator Hda (5), as well as DNA ligase (2). Additionally, it facilitates DNA damage tolerance and repair pathways through interactions with specialized translesion synthesis polymerases (Pol II, Pol IV, and Pol V) (6-9), and with mismatch repair proteins (MutL and MutS) (2). Recently, β-clamp was also shown to be involved in post-replication gap repair (10).

The molecular basis for these diverse interactions lies in a conserved hydrophobic pocket present on each monomer of the homodimeric β-clamp. Partner proteins possess specific clamp-binding motifs (CBMs) that engage with these pockets (1). Studies have demonstrated that β-clamp can simultaneously accommodate two different binding partners, with each protein occupying one pocket each (11). The canonical CBM was initially characterized as QL[S/D]LF (12). However, recent analyses have led to revised consensus sequences, such as Qφx[L/M][F/L] and Qφx[L/M]x[F/L], where φ represents an aliphatic residue (leucine, isoleucine, valine, or alanine) and x denotes any residue (1). These CBMs typically span five to six residues, with distinct functional roles for specific residues. While the glutamine (Q) at the first position of the motif significantly enhances binding affinity, it is not absolutely required for interaction with β-clamp, as exemplified by the clamp loader’s δ subunit, which contains an alanine at this position (1). Conversely, the residues at the end of the motif are critical for binding, with the leucine-phenylalanine (LF) combination providing highest affinity (1).

The mammalian homolog of β-clamp, proliferating cell nuclear antigen (PCNA), orchestrates similar processes in eukaryotic cells (13). PCNA has an extensive interactome, comprising over 600 potential binding partners (14), many with direct interactions (15), making it a prime target for developing sensitizing and antimutagenic inhibitors in cancer treatment (16). Despite divergent quaternary structures—PCNA forming a homotrimer of two-domain monomers versus a homodimer of three-domain monomers of β-clamp—both proteins maintain a conserved hexameric architecture. This arrangement provides PCNA with three hydrophobic interaction pockets (13), compared to β-clamp’s two. Notably, while the interaction network of PCNA has been extensively characterized, β-clamp’s interactome remains comparatively understudied, suggesting the existence of undiscovered binding partners and functionalities. In eukaryotes, PCNA participates in most DNA repair pathways, including homologous recombination through Rad51 interaction (17,18). In bacteria, however, β-clamp has so far not been implicated in this process.

It is well established that the RecBCD complex, comprising RecB, RecC, and RecD subunits, manages DSB repair by processing DNA ends and facilitating loading of RecA (the bacterial Rad51 homolog) after DNA damage (19,20). RecBCD exhibits remarkable specificity through recognition of Chi sites—8-nucleotide DNA motifs (5′-GCTGGTGG-3′) that regulate homologous recombination. Upon encountering a Chi site, RecBCD transitions from rapid DNA degradation of both strands to preferential degradation of the 5′ strand, while continuing to unwind and protect the 3′ strand. This transition facilitates RecA loading and promotes precise homologous recombination (21-24).

Given RecA’s homology to Rad51 and recent evidence of β-clamp involvement in single-strand gap repair through RecF interaction (10), we hypothesized a potential role of β-clamp in DSB repair. Here, we demonstrate that β-clamp interacts directly with RecBCD in *Escherichia coli* through a distinct CBM in the RecB subunit, impacting RecBCD-dependent activities.

## MATERIAL AND METHODS

### Strain construction and growth conditions

Strains and plasmids used in this study are listed in Table 1 and Table 2, respectively. Most of the strains are derived from the adenylate cyclase deficient (*cya*^-^) *E. coli* (reporter) strain BTH101, which is a commonly used host organism for investigation of protein-protein interactions using the bacterial two-hybrid system (25,26). Gene deletions or mutations were introduced into BTH101 using standard P1 transduction (27). Single deletion mutants available in the Keio collection (28) were used to knock out *recA, recB* or *recC*, while DPB271 was used as donor for the introduction of a *recD* mutation. Flp recombinase (pCP20) (29) was used to remove the *kan*-cassette from strains transformed with plasmids carrying kanamycin resistance. Variants of pKT25 and pUT18C plasmids harboring *dnaN* and *recB* (wild type and mutants) were constructed by GenScript Inc (Piscataway, NJ, USA). His-tagged *recB* was cloned into the pBAD30 vector (pBAD30-*recB*) by Dongxuan Gene Technology Co., Ltd (Guangzhou, China). Strains from frozen glycerol stock were streaked onto LB-agar plates with appropriate antibiotics and supplements for selection and incubated at 37°C, unless otherwise specified.

**Table 1.** *E. coli* strains used in this study. P1 transduction is shown as: P1 Donor x Recipient. Relevant antibiotic resistances are shown in brackets: Kan^R^, Kanamycin resistance; Str^R^, Streptomycin resistance; Tet^R^, Tetracycline resistance.

| Strain | Relevant genotype | Source |
| --- | --- | --- |
| BTH101 | F<sup>-</sup>, *cya*-99, *araD*139, *galE*15, *galK*16, *rpsL*1 (Str<sup>r</sup>), *hsdR*2, *mcrA*1, *mcrB*1 [Str<sup>R</sup>] | Euromedex BACTH system kit (30) |
| JW2788 | BW25113 Δ*recB::kan* [Kan<sup>R</sup>] | Keio collection (28) |
| IMR04 | BTH101 Δ*recB* [Str<sup>R</sup>] | This work; P1 JW2788 x BTH101 (*kan* removed with pCP20) |
| JW2790 | BW25113 Δ*recC::kan* [Kan<sup>R</sup>] | Keio collection (28) |
| IMR05 | BTH101 Δ*recC* [Str<sup>R</sup>] | This work; P1 JW2790 x BTH101 (*kan* removed with pCP20) |
| DPB271 | $\lambda^-$ *recD*1903::minitet (*recD::minitet*) [Tet<sup>R</sup>] | Biek DP and Cohen SN (31) |
| IMR07 | BTH101 *recD::minitet* [Tet<sup>R</sup>] | This work; P1 DPB271 x BTH101 |
| JW2669 | BW25113 Δ*recA::kan* [Kan<sup>R</sup>] | Keio collection (28) |
| IMR13 | BTH101 Δ*recA* [Str<sup>R</sup>] | This work; P1 JW2669 x BTH101 (*kan* removed with pCP20) |
| IMR19 | BTH101 $\Delta recA::kan \Delta recB$ [Str<sup>R</sup>, Kan<sup>R</sup>] | This work; P1 JW2669 x IMR04 |
| BHC132 | MG1655 *recB::HaloTag* | Meriem El Karoui (32) |
| OZA001 | MG1655 *gfp-dnaN frt-kan* [Kan<sup>R</sup>] | Tsutomu Katayama (33) |
| EH219 | MG1655 *gfp-dnaN recB::HaloTag* [Kan<sup>R</sup>] | This work; P1 OZA001 x BHC132 |

**Table 2.**
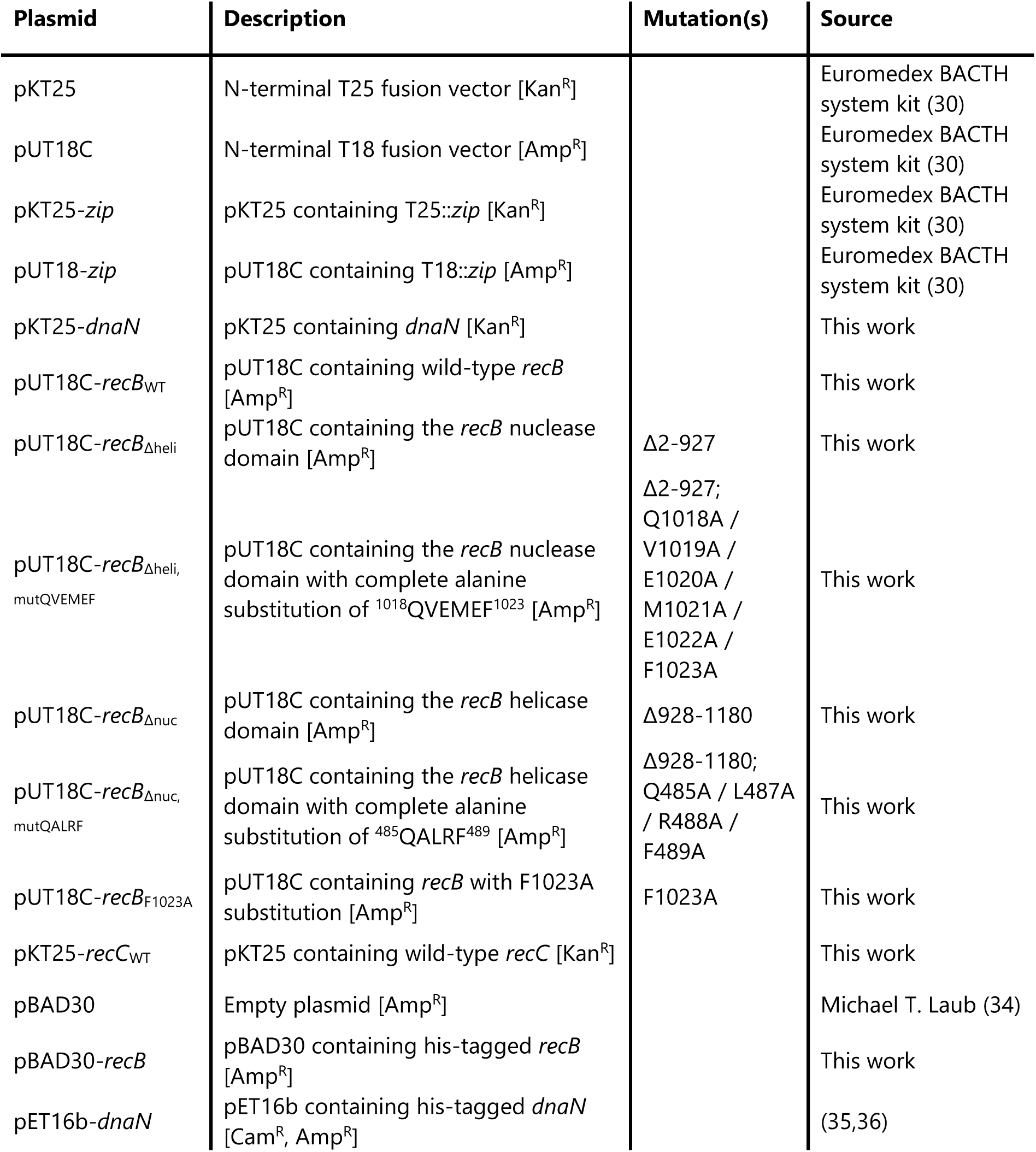
Main plasmids used in this study. Relevant antibiotic resistances are shown in brackets: Amp^R^, Ampicillin resistance; Cam^R^, Chloramphenicol resistance; Kan^R^, Kanamycin resistance.

### Yeast two-hybrid screening

Yeast-two hybrid screening was outsourced to Hybrigenics (Paris, France). This service employs a high-complexity *E. coli* prey library (∼3 x107 clones) and follows a robust mating-based screening protocol to identify binary protein interactions. Full-length *E. coli dnaN* (encoding β-clamp) was cloned into the Gal4 DNA-binding domain expression vector pGBKT7, which carries LEU2 and TRP1 selection markers (bait plasmid). The *dnaN* bait was screened against an *E. coli* cDNA/genomic prey library provided by Hybrigenics, totaling approx. 29 million interaction tests. Transformants were selected on medium lacking leucine, tryptophan, and histidine, supplemented with 0.5 mM of 3-amino-1,2,4-triazole (3-AT) to suppress background growth. Positive interactions were confirmed and sequenced by Hybrigenics.

### Bacterial two-hybrid (BACTH) assay

Protein-protein interactions between RecB and β-clamp were examined using the bacterial adenylate cyclase-based two-hybrid (BACTH) system (Euromedex), as previously described (37), with optimizations noted below. This system relies on the functional reconstitution of *Bordetella pertussis* adenylate cyclase (CyaA) fragments T25 and T18, which are genetically fused to the proteins of interest. Upon interaction of the fusion partners, cAMP production is restored in the *cyaA*-deficient *E. coli* BTH101 strain, activating downstream reporters such as *lacZ* and *mal* operons. All co-transformants were tested using three complementary readouts: (1) qualitative X-gal spot assay, (2) quantitative β-galactosidase assay, and (3) semi-quantitative growth in M63 minimal medium with maltose.

#### Strain Cultivation and Plasmid Transformation

Recipient strains (BTH101 or derivatives) were streaked from glycerol stock onto LB agar plates containing streptomycin (100 µg/mL), Isopropyl β-D-1-thiogalactopyranoside (IPTG) (0.5 mM), and 5-bromo-4-chloro-3-indolyl β-D-galactopyranoside (X-gal) (40 µg/mL) and incubated at 37°C until cya⁺ (blue) revertants became visible among the cya⁻ (white) colonies. Multiple white colonies were picked and cultured overnight in LB with streptomycin (100 µg/mL) at 37°C. These cultures were diluted 1:50 and expanded before electroporation.

Plasmid constructs (typically ∼30 ng DNA) were introduced by electroporation into electrocompetent cells, which were prepared by growing to optical density at 600 nm (OD_600_) around 0.5, followed by three washes in ice-cold ultrapure water (Millipore, Milli-Q) and final resuspension in 40 µL ice-cold Milli-Q. Electroporation was carried out in 1 mm gap cuvettes (Thermo Fisher Scientific Inc., Waltham, MA, USA) at 1.35 kV, 600 Ω, and 10 µF using a Gene Pulser II (Bio-Rad). Transformed cells were immediately recovered in 1 mL SOC at 37°C for 1.5 h with shaking and plated on LB agar supplemented with appropriate antibiotics (Table 3). Plates were incubated at 30°C for up to 3 days.

**Table 3.** Antibiotics and inducers used in BACTH experiments. * Used when required for selection of specific constructs.

| Additive | LB medium | M63 minimal medium |
| --- | --- | --- |
| Streptomycin | 100 $\mu\text{g/mL}$ | 50 $\mu\text{g/mL}$ |
| Kanamycin | 50 $\mu\text{g/mL}$ (plasmid); 30 $\mu\text{g/mL}$ (strain) | 25 $\mu\text{g/mL}$ (plasmid) |
| Ampicillin | 100 $\mu\text{g/mL}$ | 50 $\mu\text{g/mL}$ |
| Tetracycline | 5 µg/mL* | 2.5 µg/mL* |
| IPTG | 0.5 mM | 0.5 mM |
| X-gal | 40 µg/mL | - |

#### Culture Conditions for Assays

Following selection, 3-5 colonies from each transformation plate were pooled to inoculate 1 mL LB containing IPTG (0.5 mM) and the appropriate antibiotics. Cultures were incubated overnight at 30°C with shaking. The same overnight culture was used to seed all subsequent assays. The following day, cultures were diluted 1:100 into fresh LB (containing IPTG and antibiotics) and grown at 30°C to mid-exponential phase (OD_600_ around 0.5-0.6, measured by Nanodrop One, Thermo Fisher Scientific Inc.) before use in assays.

#### Assay I: X-gal Spot Assay (Qualitative)

The X-gal spot assay was used to visualize protein-protein interactions via blue/white colorimetric readout. Mid-log cultures were serially diluted (10-fold series), and 5 µL of each dilution (including undiluted) was spotted onto LB agar containing IPTG (0.5 mM), X-gal (40 µg/mL), and selective antibiotics. To assess interaction stability under genotoxic stress, an identical plate was exposed to UV light (15 J/m²). Plates were incubated at 30°C for 2-3 days, and images were captured with a smartphone camera.

#### Assay II: β-galactosidase Activity Assay (Quantitative)

β-galactosidase activity was quantified using a modified Miller assay adapted for 96-well format. Cultures at OD_600_ around 0.5-0.6 were measured again in a plate reader (Victor Nivo, PerkinElmer, Waltham, MA, USA) to account for instrument variability. Cells were pelleted, resuspended in PM2 buffer (70 mM Na₂HPO₄, 30 mM NaH₂PO₄, 1 mM MgSO₄, 0.2 mM MnSO₄, and 100 mM β-mercaptoethanol), and permeabilized with 0.01% (w/v) SDS and 10% (v/v) toluene for 45 min at 37°C. Supernatants were transferred in triplicate to 96-well polypropylene plates and pre-equilibrated at 30°C. The substrate o-nitrophenyl-β-D-galactoside (ONPG) was added to a final concentration of 0.67 mg/mL. After 30-45 min, the reaction was stopped with 200 mM Na₂CO₃. Absorbance at 405 nm was recorded, and β-galactosidase activity calculated in Miller Units using the following equation:

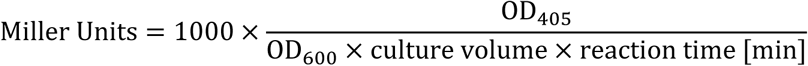

#### Assay III: Selective Growth in M63 Minimal Medium with Maltose (Semi-Quantitative)

This assay exploits the cAMP dependence of the *mal* regulon, allowing only cya^+^ cells with interacting fusion proteins to grow on maltose as sole carbon source. Cultures were first adjusted to an OD_600_ of 1 in LB and then diluted 1:100 into M63 minimal medium (2 g/L (NH₄)₂SO₄, 13.6 g/L KH₂PO₄, 0.5 mg/L FeSO₄·7H₂O, 1 g/L thiamine), supplemented with 0.4% (w/v) maltose, 0.5 mM IPTG, and appropriate antibiotics. Cultures were incubated in triplicate at 30°C with shaking, and OD_600_ was measured at 24, 48, and 72 hours using a plate reader. A negative control (M63-Mal without cells) was included to monitor contamination. Growth curves were visualized in GraphPad Prism (Dotmatics).

### Preparation of cell lysate for co-immunoprecipitation

His-tagged *recB* was cloned into the pBAD30 plasmid (kind gift from Prof. Michael T. Laub, MIT and Howard Hughes Medical Institute, USA) by Dongxuan Gene Technology Co. and transformed into *E. coli* MG1655. An overnight culture was diluted 1:100 into 1 L of LB broth supplemented with ampicillin (100 µg/mL). When the culture reached an OD₆₀₀ of 0.5, expression was induced with 0.4% (w/v) L-arabinose. After 2 hours of induction at 37°C, cells were harvested by centrifugation (4000 x *g*, 10 min), washed with PBS, and crosslinked with 1% formaldehyde for 20 min at room temperature. Crosslinking was quenched by the addition of glycine to a final concentration of 2.5 M. The cell pellets were washed three times with PBS, flash-frozen in liquid nitrogen, and stored at −80°C.

For lysis, cell pellets were resuspended in 10 mL of lysis buffer (50 mM Tris-HCl, pH 8.0; 500 mM NaCl; 10 mM β-mercaptoethanol; 1 mg/mL lysozyme) and subjected to sonication in 3 × 15 seconds pulses (Vibra cell, Sonics & Materials, Inc., Newtown, CT, USA). The lysate was clarified by centrifugation at 20,000 × *g* for 20 min at 4°C, and the supernatant was collected for subsequent analysis.

### Co-immunoprecipitation

Co-immunoprecipitation was performed using Dynabeads His-Tag Isolation and Pulldown beads (Invitrogen, Thermo Fisher Scientific Inc.), following the manufacturer’s instructions. Briefly, 2 mg of total protein from the clarified lysate was diluted in 700 µL of pull-down buffer (3.25 mM sodium phosphate, pH 7.4; 70 mM NaCl; 0.01% Tween-20) and incubated with 1 mg (25 µL) of the His-tag beads. As a negative control, Dynabeads Protein A (Invitrogen) were incubated with 1 µg of mouse polyclonal IgG (Diagenode, Liège, Belgium, C15400001-15) and the same lysate.

After incubation for 30 min at 4°C with gentle rotation, beads were washed four times with wash buffer (50 mM sodium phosphate, pH 8.0; 300 mM NaCl; 0.01% (v/w) Tween-20). Proteins were eluted by incubation in 50 µL of elution buffer (wash buffer supplemented with 300 mM imidazole) for 5 min at room temperature. Beads were removed magnetically, and eluates were analyzed by SDS-PAGE. Wild-type his-tagged β-clamp and RecB was expressed from pET16b-*dnaN* and pBAD30-*recB*, respectively, and purified with Protino Ni-NTA Agarose (AH Diagnostics/MACHEREY-NAGEL, Düren, Germany) for use as controls.

### Imaging and analysis of RecB and β-clamp colocalization

Fixed cells with endogenously tagged *recB* and *dnaN* (strain EH219) were imaged. β-clamp was N-terminally Green fluorescence protein (GFP)-tagged (33) (kind gift from Tsutomu Katayama), while RecB had a HaloTag inserted in a loop after residue S47 (32) (kind gift from Meriem El Karoui), allowing staining with Janelia Fluor 549 HaloTag ligand (JF549-HTL) (Promega Corporation, no. GA1110, Madison, WI, USA).

Strains were first grown overnight on LB-agar plates with kanamycin for selection. Then, 2 mL LB with kanamycin was inoculated with 5 plate colonies to create overnight cultures (ONCs). The next day, ONCs were diluted 1:200 in fresh LB medium with kanamycin and cultured at 37°C until exponential growth (OD_600_ of 0.10). JF549-HTL (5 µM final concentration) was then added to the culture and incubation continued for another 60 min for staining the RecB HaloTag, before washing five times with LB. After staining, samples were taken for fixation before and after 2 and 20 min of ciprofloxacin (20 ng/mL) exposure. Samples were fixed by resuspending and incubating them in PBS with 4% (v/w) formaldehyde for 60 min, followed by washing twice with PBS and resuspension in PBS for imaging.

Imaging and analysis commenced according to the procedures previously described in Vikedal et al. (37). Briefly, we used a Nikon Eclipse Ti2-E inverted microscope equipped with a 60x oil objective, a CrestOptics X-Light V3 spinning disk confocal module (50:400 µm spinning disk), a Lumencor Celeste multi-line laser and two Teledyne Photometrics Kinetix sCMOS cameras. Fixed samples were transferred onto PBS-agar pads (PBS with 1% (w/v) agarose) on microscope slides within a Gene Frame (Thermo Scientific, no. AB0576), and the sample sealed with a cover slip (#1.5 thickness). Three locations were imaged for each sample to get enough cells for analysis. Three channels were used for fluorescence imaging: a transmitted light channel for cell outlines, in addition to GFP and JF549 fluorescence channels. GFP was excited at 477 nm with emission collected between 501-521 nm. JF549 was excited at 546 nm with emission collected between 580-610 nm. Flat-field correction was applied to all images before analyses. A beta-version of MicrobeJ (beta-version 5.13p (20)) (38) was used to segment cells and detect foci of GFP–β-clamp and RecB-HaloTag/JF549 within them. The foci numbers per cell were quantified along with their colocalization. Foci were defined as colocalized if their centers (maxima) were at most one pixel apart in any direction (pixel size is 0.108 µm).

### Peptides and proteins

All peptides were purchased from TAG Copenhagen (Søborg, Denmark). Pol III C-terminal (EQVELEFD), Pol III internal (GQADMFG), RecB nuclease motif (KQVEMEFY) and RecB nuclease motif F-A (KQVEMEAY) had one additional N- and C-terminal flanking residue outside the proposed binding motif. To increase the solubility of the RecB helicase motif peptides, one C-terminal and three flanking N-terminal residues were added to this RecB motif (GKNQALRFV) and its variant motif F-A (GKNQALRAV). All peptides, except Pol III C-terminal (EQVELEFD), were N-terminally acetylated and C-terminally amidated to remove charges in the N-and C-termini, not present in the context of the full-length proteins.

Unlabeled and 2H, 13C, 15N labelled *E. coli* β-clamp was expressed and purified as previously described (39). The β-clamp concentration was determined by absorbance at 280 nm measured on a NanoDrop ND-1000 spectrophotometer (Thermo Fisher Scientific Inc.) and using an extinction coefficient calculated by the Expasy ProtParam web tool (https://web.expasy.org/protparam/) (40).

### Nuclear Magnetic Resonance (NMR) spectroscopy

#### Saturation Transfer Difference (STD)-NMR

All STD-NMR samples contained 150 mM peptide with or without the addition of 1.5 mM unlabeled β-clamp in 20 mM sodium phosphate, 100 mM NaCl, 5 mM DTT, 0.3% (v/v) DMSO, 5% (v/v) D_2_O and 125 mM 2,2-dimethyl-2-silapentane-5-sulfonate (DSS), pH 7.4. Since most peptides did not contain any tryptophan or tyrosine residues, peptide concentrations for the STD-NMR samples were calculated based on the molecular weight of the peptides. The samples were transferred to 5 mm NMR tubes. STD NMR experiments were recorded at 25°C on a Bruker Avance III HD 600 MHz NMR spectrometer equipped with a 5mm QCI Cryoprobe. All experiments were recorded using a standard Bruker STD NMR setup with an off-resonance frequency at -40 ppm and on-resonance at 0 ppm. The number of scans was 1024.

#### 2D TROSY-^1^H-^15^N-HSQC

2D TROSY-^1^H-^15^N-HSQC (Two-Dimensional Transverse Relaxation Optimized Spectroscopy Heteronuclear Single Quantum Coherence experiment correlating ^1^H and ^15^N nuclei) spectra were recorded of 140 mM ^2^H, ^13^C, ^15^N labelled β-clamp with and without the addition of 700 mM RecB nuclease motif peptide (KQVEMEFY) in 20 mM sodium phosphate, 100 mM NaCl, 5 mM DTT, 5% (v/v) D2O, 125 mM DSS, 2% (v/v) DMSO, pH 7.4. The estimation of the RecB motif peptide stock concentration used for the NMR HSQC experiment was based on the integration of protons from the peptide compared to the integration of protons of the DSS signal, for which the concentration in the sample is known. The samples were transferred to 5 mm Shigemi tubes, and the NMR spectra recorded at 37°C on a Bruker Avance III 750 MHz spectrometer equipped with a cryoprobe. DSS was used to reference ^1^H, and ^15^N and ^13^C were referenced indirectly using their gyromagnetic radii. The spectra were processed in Topspin (Bruker) and analyzed in CcpNmr Analysis (41). The β-clamp assignment was acquired from BMRB under the deposition number 52494 (39). Chemical shift perturbations (CSPs) were calculated using the following equation (42):

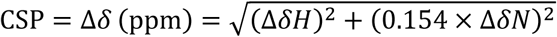

### Isothermal Titration Calorimetry (ITC)

Unlabeled β-clamp was buffer exchanged into 20 mM sodium phosphate, 100 mM NaCl, 1 mM tris (2-carboxyethyl)phosphine (TCEP), pH 7.4 using a 15 mL spin filter 10000 MWCO (Millipore, Darmstadt, Germany). DMSO was added to the β-clamp sample to a final concentration of 0.02% (v/v). RecB nuclease motif peptide (KQVEMEFY) (dissolved in 100% DMSO) was diluted into the same buffer as above, with a final DMSO concentration of 0.02% (v/v). 70.6 mM RecB nuclease motif peptide (KQVEMEFY) was added to the cell (the peptide concentration estimated by NMR as described above) and 278-294 mM β-clamp was added to the syringe. Peptide and protein samples were centrifuged at 20,000 g at 25°C for 15 min to degas prior to loading. The ITC experiments were repeated three times and were recorded at 25°C on a Malvern MicroCal PEAQ-ITC instrument (Malvern, Worcestershire, United Kingdom) using a stir speed of 750 rpm. The ITC data was fit to a one-site binding model using the MicroCal PEAQ-ITC Software. Reported errors are standard error of the mean (SEM) values of the three ITC replicates.

### Survival Assays

The *E. coli* BTH101 *ΔrecB* strain (IMR04) was transformed with pUT18C plasmids encoding either wild-type RecB or the mutant RecB_F1023A_, as well as with the empty pUT18C vector. Non-transformed BTH101 and BTH101 *ΔrecB* served as positive and negative controls, respectively. All strains were cultured at 30°C to early exponential phase (OD_600_ around 0.2, measured by Nanodrop One) before treatment.

#### UV Survival Assay

For UV exposure, bacterial cultures were serially diluted 10-fold in LB. From each dilution, 5 µL was spotted onto four separate LB agar plates supplemented with IPTG (0.5 mM) and the appropriate antibiotics. Once droplets had dried, one plate was kept unexposed as a control, while the remaining three plates were subjected to UV irradiation at doses of 5, 15, or 30 J/m² using an CL-1000 Ultraviolet crosslinker (UVP). Plates were incubated overnight at 30°C, after which colony-forming units (CFUs) were counted. Survival at each UV dose was calculated as the ratio of CFUs on the UV-exposed plates to CFUs on the unexposed control plate.

#### Ciprofloxacin Survival Assay

To assess ciprofloxacin sensitivity, bacterial cultures at OD_600_ around 0.2 were divided into five equal subcultures and treated with ciprofloxacin at final concentrations of 0, 4, 8, 12, or 16 ng/mL. Cultures were incubated at 37°C with shaking for 2 hours. Following treatment, cells were pelleted by centrifugation and resuspended in fresh LB medium to remove residual antibiotic and halt exposure. Cultures were then serially diluted 10-fold, and 5 µL of each dilution was spotted onto LB agar plates. After overnight incubation at 30°C, CFUs were counted. Survival at each ciprofloxacin concentration was calculated relative to the number of CFUs in the untreated control.

### DNA degradation assay

*E. coli* cells lacking both *recA* and *recB* were transformed with pUT18C plasmids expressing either wild-type RecB, the RecB_F1023A_ mutant, or the empty vector. IMR13 (*ΔrecA*) served as a positive control for RecB-mediated DNA degradation, while IMR19 (*ΔrecA ΔrecB*) was used as a negative control.

Overnight cultures were diluted to an OD_600_ of 0.01 into LB medium supplemented with appropriate antibiotics and incubated with shaking at 37°C until they reached an OD_600_ of 0.2. Each culture was split into two 1 mL aliquots. One aliquot was fixed immediately in ethanol (50% final concentration) to assess DNA content prior to UV exposure. The second aliquot was pelleted, resuspended in 1×PBS, and exposed to UV light at a dose of 50 J/m² (CL-1000 Ultraviolet crosslinker, UVP). Immediately after UV exposure, cells were pelleted again, resuspended in fresh LB (with antibiotics), and incubated at 37°C with shaking for 3 hours to allow DNA degradation to proceed. Following the post-UV incubation, cultures were fixed in ethanol (50% final concentration) and stored at 4°C until further processing.

#### Microscopy

For imaging of DNA content, cells were stained with 5 µg/mL Hoechst 33258 in PBS for 10 min at room temperature. Following staining, samples were washed three times with PBS to remove excess dye. To normalize for differences in cell density across samples, cell suspensions were appropriately concentrated to ensure comparable and sufficient cell numbers for imaging. For microscopy, 10 µL of each cell suspension was applied to agarose pads (1% [w/v] agarose in PBS) prepared within a Gene Frame (Thermo Fisher Scientific Inc., Cat. No. AB0576) mounted on a standard microscope slide. After drying, pads were sealed with a #1.5 thickness coverslip.

Imaging was performed on a Leica DM6000 B fluorescence microscope equipped with a 100x oil immersion objective (Leica HCX Plan Apochromat 100x, 1.40 NA, PH3 CS), a Leica EL6000 metal-halide illumination system, and a Hamamatsu C9100-14 EM-CCD camera. For each sample, six fields of view were acquired, each including both phase-contrast and fluorescence channels. Fluorescence imaging was conducted using a narrow band-pass filter set optimized for CFP (Excitation: 426-446 nm; Emission: 460-500 nm). All imaging parameters—including exposure time, illumination intensity, and gain—were kept constant across all strains, treatments, and replicates to ensure quantitative comparability and minimize experimental bias.

#### Quantification of DNA Degradation

Microscopy images were analyzed using the open-source software Fiji (ImageJ) (43), along with the MicrobeJ plugin (version 5.13l) (38). Due to minor misalignment between the phase-contrast and fluorescence channels in some images, the *Channel Aligner* tool in MicrobeJ was used to adjust alignment prior to analysis. The phase-contrast channel was then used for cell detection and segmentation using the “Medial Axis” method. Detection parameters were constrained based on cell morphology, including area, length, width, circularity, curvature, sinuosity, angularity, and intensity.

To assess DNA content, the mean fluorescence intensity of each segmented cell was measured in the Hoechst (CFP) channel across six acquired images per sample. Background correction was applied by measuring the mean fluorescence intensity from cell-free regions in each image and subtracting this value from the corresponding cellular fluorescence intensities.

### Western blot

Samples, including purified RecB, β-clamp, input (10 µg; 0.5%), and pull-down eluates, and samples from BTH101 cultures used in survival assays, were mixed with 1 mM dithiothreitol (DTT) and 1×NuPAGE LDS sample buffer, then heated to 75°C for 10 min. Proteins were resolved on 4-15% Mini-PROTEAN TGX precast gels (Bio-Rad) using Tris/Glycine/SDS running buffer. SeeBlue Plus2 Pre-stained Protein Standard (Invitrogen) was used as a molecular weight marker.

Proteins were transferred onto 0.2 µm nitrocellulose membranes using the Trans-Blot Turbo Transfer System (Bio-Rad), and membranes were blocked in 5% (w/v) skim milk (Merck) in PBS-T (PBS with 0.1% Tween-20) for 45-60 min at room temperature. Membranes showing RecB levels in BTH101 cultures were additionally blocked with cell extracts from *ΔrecB* cells. These cell extracts were made as previously explained (37) with some alterations; anti-RecB polyclonal antibody used (rabbit, MyBiosource, cat. no. MBS7162389) and no ciprofloxacin added.

Membranes were incubated overnight at 4°C with primary antibodies diluted in blocking buffer: anti-RecB at 1:200, and anti-β-clamp antibody (rabbit, Eurogentec, Seraing, Liège, Belgium) at 1:20,000. After washing, membranes were incubated for 45 min with goat anti-rabbit IgG (H+L), HRP-conjugated secondary antibody (MyBiosource, cat. no. MBS705310), followed by detection using SuperSignal West Femto Maximum Sensitivity Substrate (Thermo Fisher Scientific, cat. no. 34577). Chemiluminescent signals were visualized using the ChemiDoc MP Imaging System and Image Lab software (Bio-Rad).

### Statistical analysis

Quantitative data were analyzed and plotted using GraphPad Prism (version 10.4.1). Differences in β-galactosidase activities were analyzed using ordinary one-way ANOVA with Dunnett correction for multiple comparisons, while DNA degradation comparisons were made using ordinary one-way ANOVA with Šidák correction. Comparisons of RecB–β-clamp colocalization before and after ciprofloxacin exposure were analyzed using repeated measures one-way ANOVA with Tukey correction. Figures are color-coded to visually indicate significant differences between tested strains, while asterisks denote the level of statistical significance. Two-tailed P-values below 0.05 were considered significant and indicated with asterisks as follows: \**P* ≤ 0.05; \*\**P* ≤ 0.01; \*\*\**P* ≤ 0.001; \*\*\*\**P* ≤ 0.0001.

## RESULTS

### RecBCD interacts with β-clamp

Given the high structural conservation between bacterial and eukaryotic sliding clamps (12), and the large number of interaction partners of the eukaryotic sliding clamps (14), it is plausible that many partners of β-clamp (DnaN) remain to be discovered. The sliding clamp has proven essential for multiple repair mechanisms in eukaryotes, including repair by homologous recombination. An initial yeast two-hybrid screening with β-clamp as bait indicated a possible interaction between β-clamp and the RecB subunit of the RecBCD repair complex (Supplementary Table S1). We, therefore, further explored this possible interaction to decipher whether β-clamp is also involved in homologous recombination in bacteria.

We employed a bacterial two-hybrid (BACTH) system to verify the RecB–β-clamp interaction from the yeast two-hybrid screening (25,26). The *recB* gene was cloned into a pUT18C vector, while *dnaN* was cloned into a pKT25 vector. These constructs were co-transformed into the *E. coli* reporter strain BTH101, which lacks endogenous adenylate cyclase (*cyaA*). In addition, the constructs were co-transformed into BTH101 variant strains with single-gene deletions of *recA*, *recB*, *recC*, or *recD* to assess the interaction’s reliance on RecA or RecBCD complex formation.

We observed a distinct interaction between RecB and β-clamp using the BACTH assay, as indicated by elevated β-galactosidase activity after 24 hours in both wild-type and *ΔrecB* BTH101 backgrounds, relative to the empty vector negative control strain (Fig. 1A). The strength of the interaction was comparable to the RecB-RecC positive control interaction (Fig. 1A). Notably, the interaction between RecB and β-clamp was abolished in BTH101 strains lacking either RecC or RecD, suggesting that the RecBCD complex interacts with β-clamp as a functional unit. The interaction initially appeared absent in the *ΔrecA* strain (Fig. 1A). However, when using an M63/maltose-based variant of the assay—which is less sensitive to differences in bacterial growth rates and allows detection of weaker or delayed interactions— the interaction was apparent also in the *ΔrecA* strain (Supplementary Fig. S1). This indicates that the RecB–β-clamp interaction likely persists in the absence of RecA, but that the slower growth rate of the *ΔrecA* strain necessitates longer incubation to achieve a detectable interaction (Supplementary Fig. S1).

**Figure 1.**
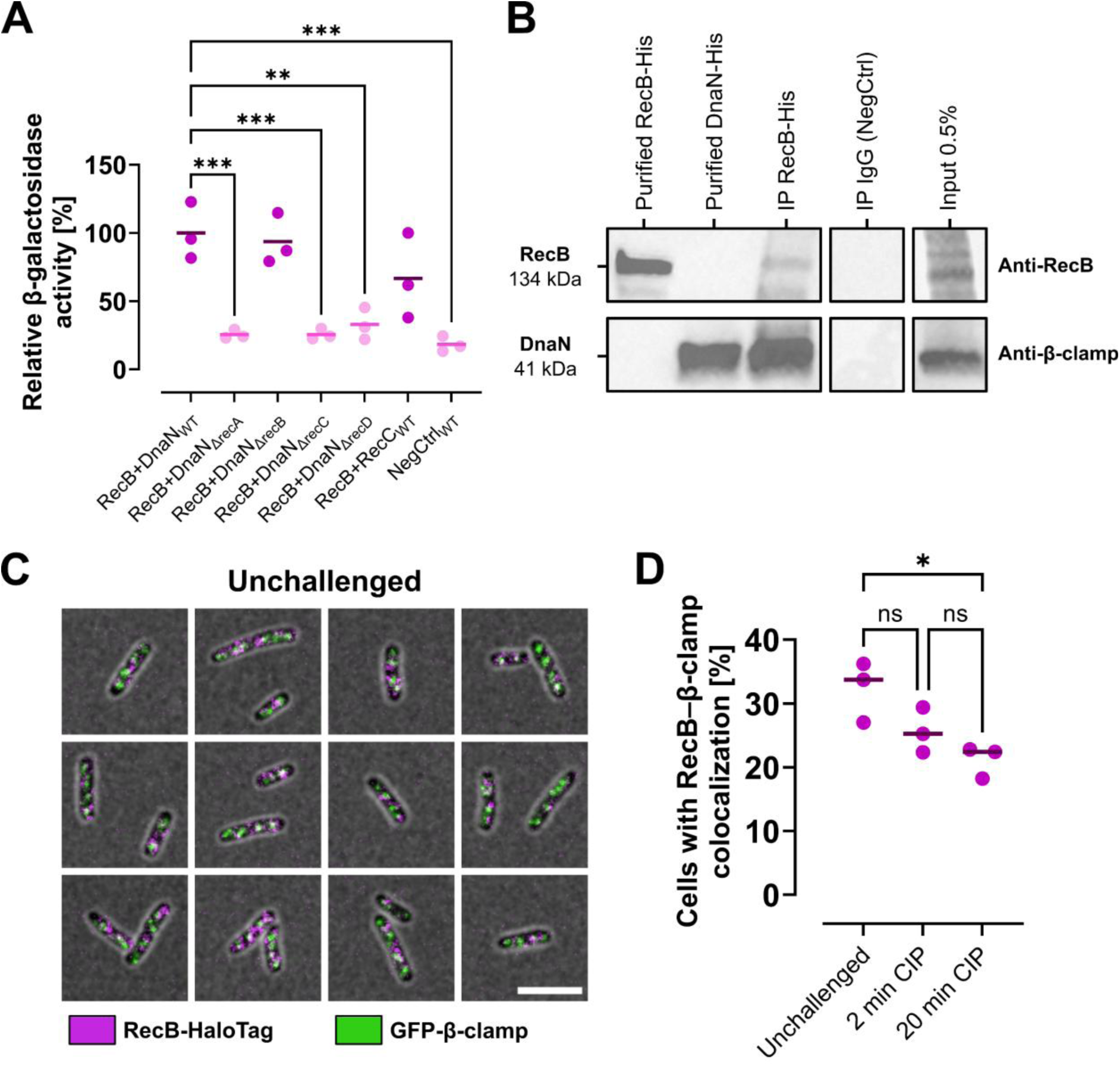
Experimental evidence for interaction between RecB and β-clamp. (**A**) Relative β-galactosidase activity from bacterial two-hybrid assay with co-expressed RecB–T18 and β-clamp (DnaN)–T25 fusion proteins in BTH101 wild type compared to *ΔrecA*, *ΔrecB*, *ΔrecC*, and *ΔrecD* deletion strains. Activities were normalized to the wild-type mean. Wild-type cells co-expressing RecB and RecC served as positive controls; negative controls used empty vectors. Strains with significantly different activity from RecB–β-clamp-expressing wild type appear in light magenta; others in dark magenta. Only significant differences are annotated. (**B**) Western blots from a co-immunoprecipitation experiment where β-clamp (DnaN) was pulled down (lower panel, anti-β-clamp) together with his-tagged RecB expressed from a pBAD30 plasmid (upper panel, anti-RecB) in an MG1655 background. (**C**) Representative images of fixed unchallenged EH219 cells displaying fluorescence from JF549-labeled RecB-HaloTag (magenta) and endogenously expressed GFP-β-clamp (green); overlapping signal appears white. (**D**) Percentage of EH219 cells showing RecB–β-clamp colocalization when unchallenged, and after 2 and 20 min of exposure to 20 ng/mL ciprofloxacin (CIP). Foci were considered colocalizing if they were at most one pixel apart in any direction (pixel size is 0.108 µm). Comparisons were made using (A) ordinary one-way ANOVA with Dunnett correction, and (D) repeated measures one-way ANOVA with Tukey correction. Lines represent means from three biological replicates; dots indicate individual replicates. ns, non-significant; \**P* ≤ 0.05; \*\**P* ≤ 0.01; \*\*\**P* ≤ 0.001.

To further validate the RecB–β-clamp interaction we performed co-immunoprecipitation experiments using a N-terminally his-tagged (6x histidine residues) version of RecB as bait, expressed from a pBAD30 vector. Indeed, β-clamp was pulled down and exhibited a strong band on the membrane (Fig. 1B).

Fluorescence microscopy was used to examine RecB localization relative to β-clamp in a strain expressing endogenous GFP-tagged β-clamp (*gfp-dnaN)* (33) and RecB with a HaloTag (32). After growth to exponential phase, RecB-HaloTag was labeled with JF549-HTL dye. One unchallenged sample was fixed, while the remaining culture was treated with ciprofloxacin at the minimum inhibitory concentration (20 ng/mL) to induce DNA DSBs. Samples were fixed after 2 and 20 min of exposure to assess RecB–β-clamp colocalization during the DNA damage response. RecB exhibited weaker foci compared to GFP-β-clamp, along with a diffuse cytoplasmic signal. A recent publication suggests that persistent RecB foci likely represent DNA-bound RecB complexes (44). Accordingly, images were processed to reduce the diffuse background while preserving the clear RecB foci (representative cells shown in Fig. 1C).

Quantitative colocalization analysis showed that approximately 30% of cells exhibited RecB and β-clamp foci within a 1-pixel distance (0.108 µm) of each other under unchallenged conditions (Fig. 1D), supporting the possibility of a direct or proximal interaction. Under these conditions, β-clamp is known to primarily localize to active replication forks and also lags behind the replication forks on newly synthesized sister chromatids due to delayed unloading (45). RecBCD plays a well-documented role in maintaining faithful DNA replication in situations where the replication forks stall or collapse at endogenous obstacles, such as secondary DNA structures or tightly bound proteins (46,47). In *E. coli*, fork stalling is estimated to occur up to 5 times per replication cycle (reviewed by Michel & Sandler (48)) and has been reported to result in spontaneous DSBs in up to 18% of cells at each generation (49). Under normal growth conditions, RecBCD may therefore localize near replication forks either in a standby state or actively engaged in resolving impediments to fork progression.

After 20 min of ciprofloxacin treatment, the fraction of cells showing RecB–β-clamp colocalization decreased (Fig. 1D and Supplementary Fig. S2A). This reduction coincided with a marked decrease in the average number of β-clamp foci per cell (Supplementary Fig. S2B), likely reflecting replisome disassembly caused by the roadblock effect of ciprofloxacin-induced DNA lesions (50). The remaining β-clamp foci after ciprofloxacin treatment may reflect clamps retained on the DNA after replisome disassembly. The number of RecB foci remained largely unchanged, in accordance with previous reports (51,52) (Supplementary Fig. S2C). The observed reduction in colocalization can therefore not be used to determine the context in which interaction is most functionally relevant—whether during normal replication or in response to DNA double-strand breaks, or both. Taken together, our results from the BACTH assay, co-immunoprecipitation, and fluorescence microscopy provide compelling evidence for a previously uncharacterized interaction between the RecBCD complex and β-clamp.

### Identification of the QVEMEF motif in RecB as a functional clamp-binding motif

To further characterize the RecB–β-clamp interaction, we searched RecB’s amino acid sequence for potential CBMs. We uncovered two candidate motifs: ^485^QALRF^489^ in the helicase domain, and ^1018^QVEMEF^1023^ in the nuclease domain. These motifs share significant homology with known DNA polymerase III (Pol III) motifs (53), differing by only two or one amino acid from the Pol III CBMs QADMF and QVELEF, respectively (Fig. 2A). Investigation into the localization of the two motifs in the protein complex revealed that while ^485^QALRF^489^ is surface-exposed and readily accessible, ^1018^QVEMEF^1023^ is positioned internally near the RecC protein interface in a tunnel through which the DNA protrudes, potentially affecting its availability for β-clamp binding (Fig. 2B).

**Figure 2.**
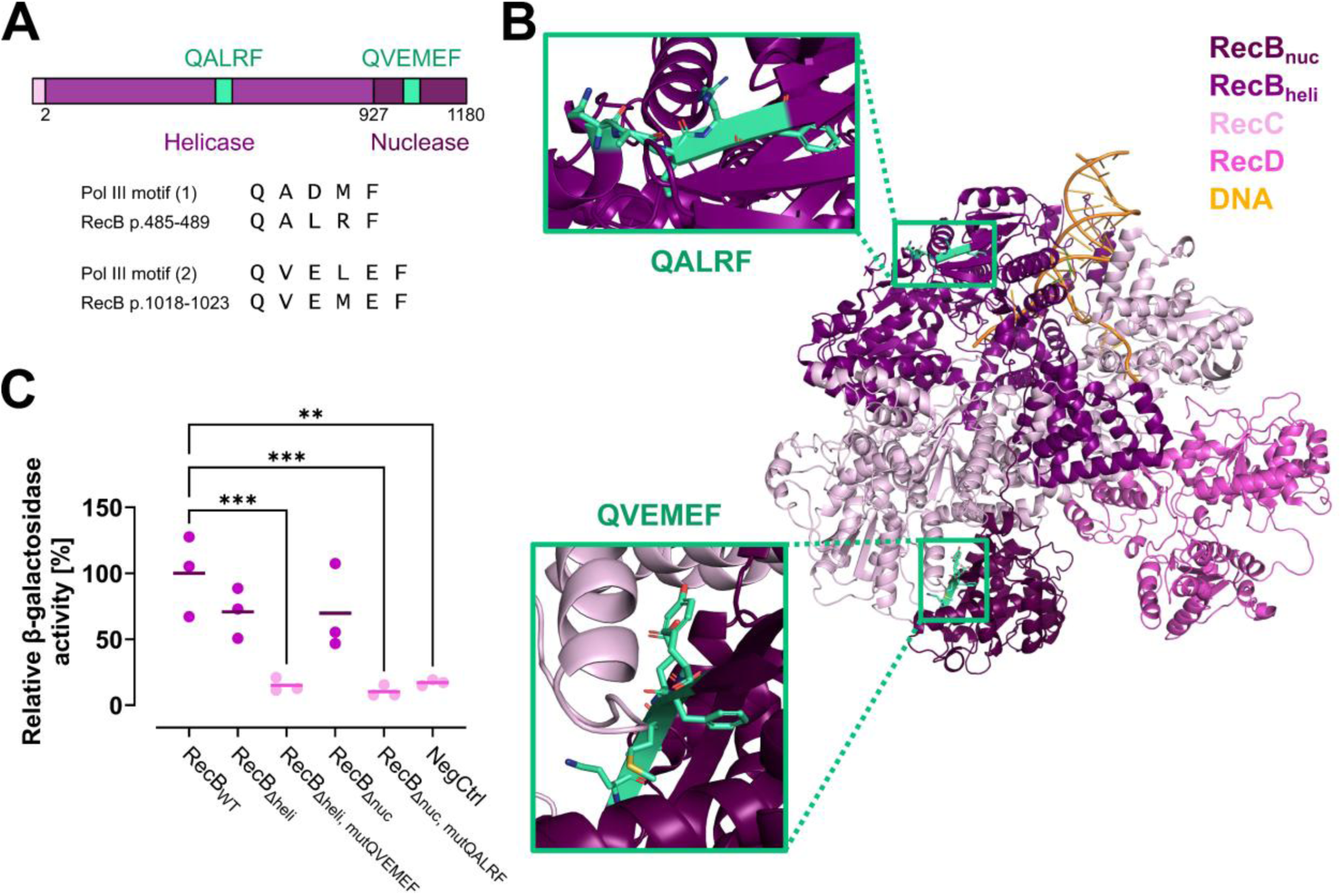
RecB may possess one clamp-binding motif (CBM) in each domain. (**A**) Sequence alignment showing two candidate CBMs (green) in RecB: QALRF (residues 485-489) in the helicase domain and QVEMEF (residues 1018-1023) in the nuclease domain. Both motifs exhibit sequence similarity to known CBMs from DNA polymerase III. (**B**) Structural mapping of the candidate motifs (green) within the RecBCD complex. QALRF is surface-exposed; QVEMEF is positioned near the RecC interface within a tunnel structure. Structure rendered from PDB ID 1W36 using PyMOL (by Schrödinger). (**C**) Relative β-galactosidase activity from bacterial two-hybrid assay of BTH101 wild type co-expressing β-clamp–T25 and various RecB–T18 variants—full-length, truncated, and mutated. RecBΔnuc includes residues 1-927; RecBΔheli includes residues 928-1180. mutQALRF and mutQVEMEF indicate complete alanine substitutions of candidate motifs. Negative controls used empty vectors. Strains with significantly different activity from full-length RecB–β-clamp-expressing wild type appear in light magenta; others in dark magenta. Only significant differences are annotated. Comparisons were made using ordinary one-way ANOVA with Dunnett correction. Lines represent means from three biological replicates; dots indicate individual replicates. \*\**P* ≤ 0.01; \*\*\**P* ≤ 0.001.

To evaluate whether our predicted CBMs in the RecB helicase and nuclease domains are functional, we constructed two truncated variants of *recB* in the pUT18C vector. The first encodes RecB residues 928-1180, thereby lacking the helicase domain of RecB, and is referred to as RecB_Δheli_. The second encodes residues 1-927, excluding the nuclease domain, and is designated RecB_Δnuc_. Using the BACTH assay, we found that both truncated variants— RecB_Δheli_ and RecB_Δnuc_—retained interaction with β-clamp (Fig. 2C). These results suggest that at least one functional CBM may be present in each domain.

To assess the contribution of specific candidate motifs, we substituted each motif entirely with alanines. In RecB_Δnuc_, the candidate QALRF motif was replaced with alanines (mutQALRF), while in RecB_Δheli_, the QVEMEF motif was similarly mutated (mutQVEMEF). These domain-specific mutations were introduced into truncated RecB variants to test each motif independently, based on the rationale that potential effects might be masked in the full-length protein due to functional compensation by the other motif.

Strikingly, BACTH analysis showed that the interaction with β-clamp was abolished in both RecB_Δnuc_ mutQALRF and RecB_Δheli_ mutQVEMEF constructs, indicating that β-clamp can interact with each domain of RecB through the identified CBMs (Fig. 2C). Alternatively, the mutations significantly affect protein folding and stability, thereby abolishing interaction indirectly (see Discussion).

To further characterize the putative CBMs in RecB, we performed nuclear magnetic resonance (NMR) experiments using recombinantly expressed and purified β-clamp and synthetic peptides representing the candidate CBMs. Each peptide included the core motif flanked by one or several native residues to mimic its native sequence context. Two peptides derived from the established CBMs of *E. coli* Pol III, QVELEF and QADMF, were included as positive controls. Samples containing peptides, but no β-clamp, were used as negative controls (Supplementary Fig. S3).

Saturation transfer difference (STD) NMR was used to assess binding interactions, where a difference in peak intensities between on-resonance and off-resonance spectra indicates ligand binding. As expected, both Pol III-derived peptides showed clear binding to β-clamp (Fig. 3A and B). Notably, the RecB-derived QVEMEF peptide also showed binding (Fig. 3C), supporting its role as a functional CBM. In contrast, a mutated QVEMEF variant in which the terminal phenylalanine was substituted with alanine (QVEMEA) showed no detectable binding (Fig. 3D), underscoring the critical role of this conserved aromatic residue in β-clamp interaction.

**Figure 3.**
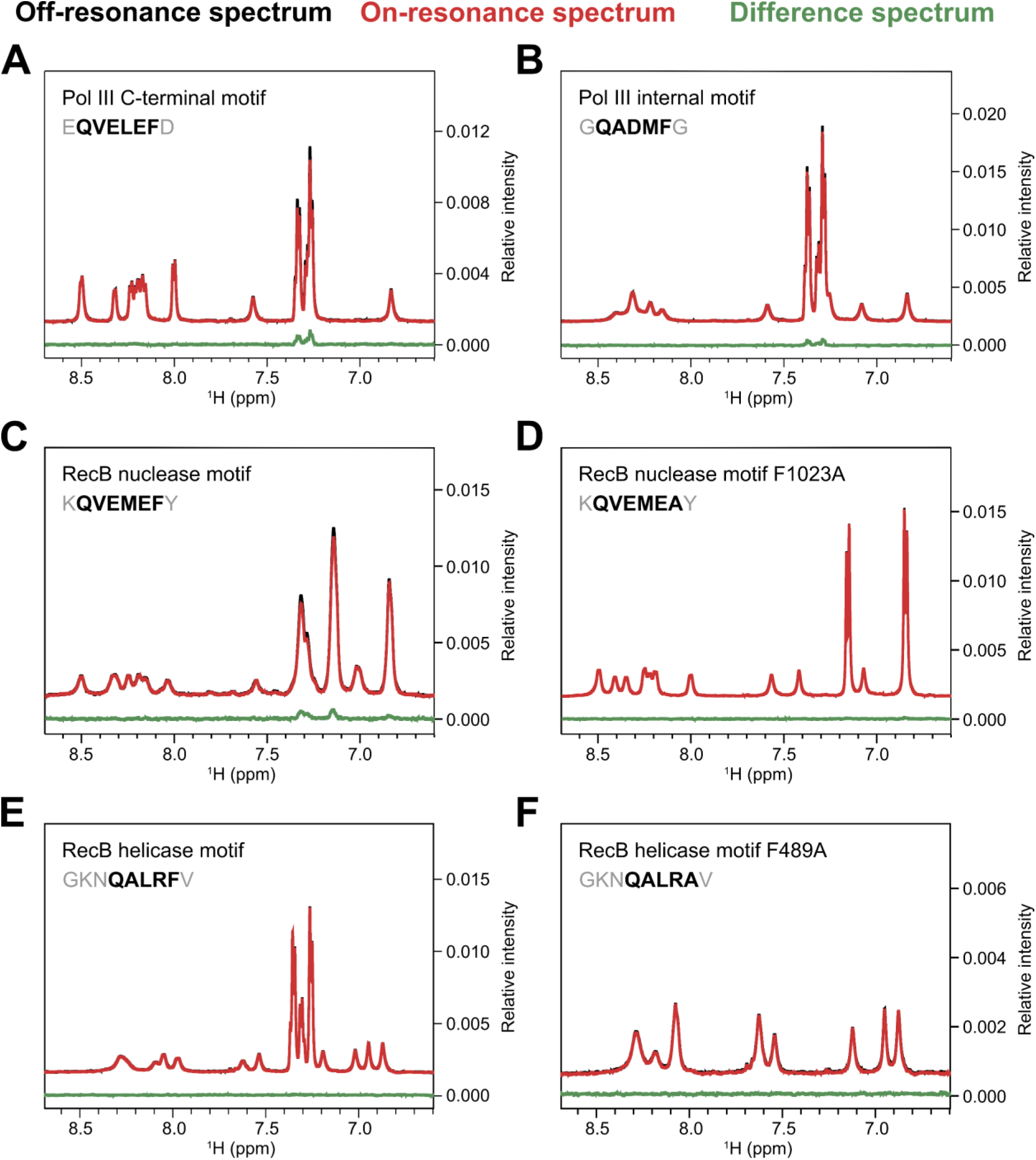
NMR analysis of β-clamp interactions with candidate clamp-binding motifs from RecB. (**A-F**) STD-NMR spectra of synthetic peptides incubated with 1% (molar ratio) purified β-clamp protein. For each sample, the off-resonance (reference) spectrum is shown in black, the on-resonance (saturated) spectrum in red, and the STD spectrum in green. Peaks in the difference spectrum indicate transfer of saturation from β-clamp to the peptide, consistent with binding. (A) QVELEF and (B) QADMF peptides from DNA polymerase III show clear binding and serve as positive controls. (C) QVEMEF peptide derived from RecB shows a distinct STD signal, indicating specific interaction. (D) A QVEMEF variant in which the terminal phenylalanine was replaced with alanine (QVEMEA) showed no detectable binding. (E) QALRF peptide from RecB and (F) its phenylalanine-to-alanine variant (QALRA) did not exhibit detectable STD signals under the conditions tested. Peptide-only controls (no β-clamp) are shown in Supplementary Fig. S3. NMR, nuclear magnetic resonance; STD, saturation transfer difference.

We also tested the second RecB motif, QALRF, and a corresponding F-to-A variant. Neither peptide showed detectable binding to β-clamp under the conditions tested (Fig. 3E and F, respectively). Although the QALRF motif is more surface-exposed in the RecBCD complex than the QVEMEF motif (Fig. 2B), its sequence deviates from the canonical CBM consensus and may therefore lack features necessary for interaction with β-clamp (1).

To quantify the binding affinity of the QVEMEF motif, we performed isothermal titration calorimetry (ITC). The interaction yielded a dissociation constant (K_D_) of 2.0 ± 0.4 µM (Fig. 4A), comparable to affinities reported for other canonical CBMs (1). Accurate determination of peptide concentration was complicated by the absence of aromatic residues, leading to an N-value below 1. However, since β-clamp was titrated from the syringe into the peptide solution in the sample cell, inaccuracies in peptide concentration affect only the N-value and not the calculated K_D_. A reduced N-value may also result from oxidation of methionine residues within the β-clamp binding pocket, previously shown to impair binding of Pol III-derived peptides (39).

**Figure 4.**
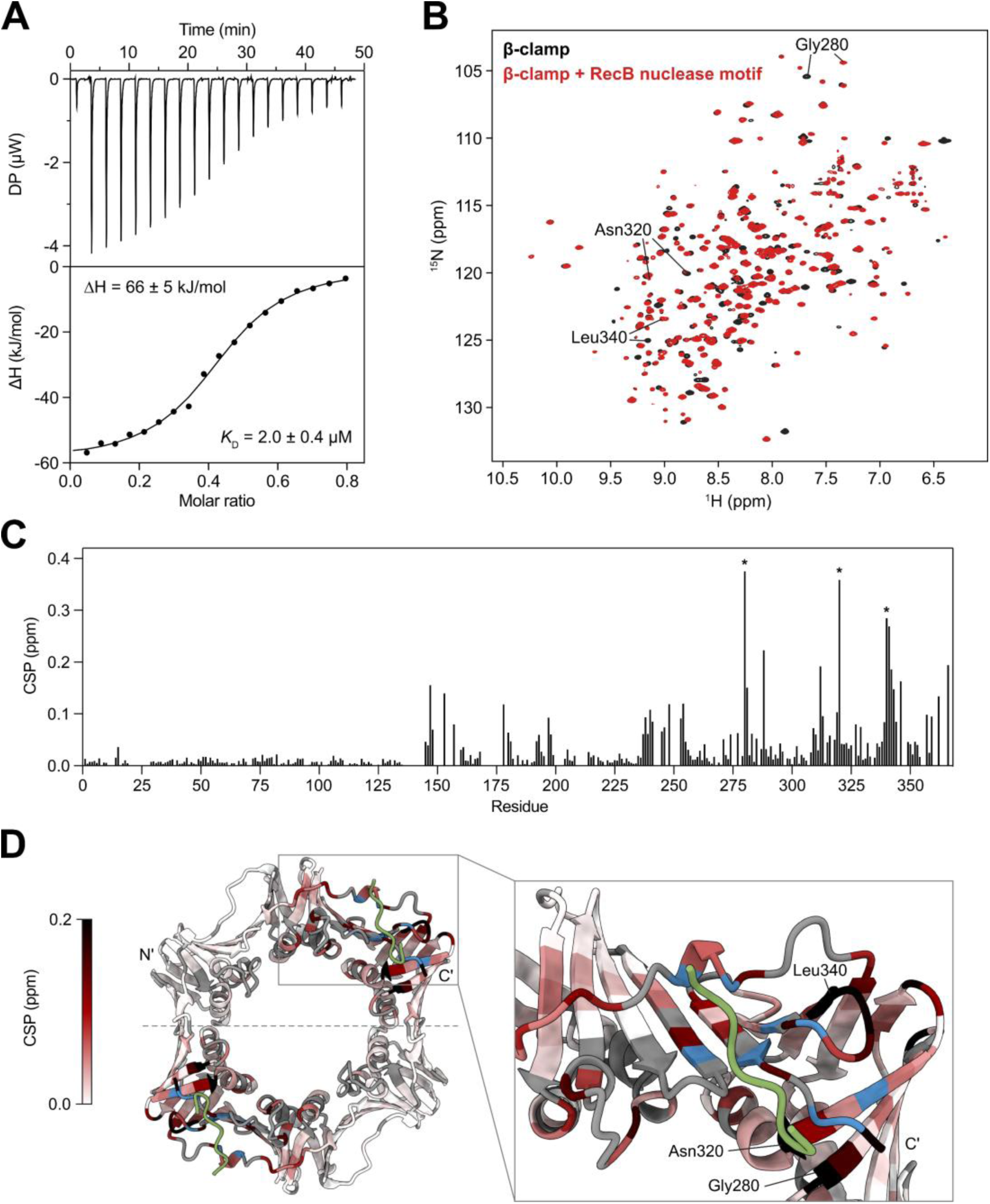
QVEMEF binds β-clamp with micromolar affinity at the canonical binding site. (**A**) ITC analysis of the interaction between the RecB-derived QVEMEF peptide and β-clamp. The upper panel shows baseline-corrected raw heat changes upon titration of β-clamp into the peptide solution; the lower panel displays the integrated binding isotherm fitted using a one-site binding model. The dissociation constant (KD) and enthalpy change (ΔH) represent the mean of three replicates; error bars indicate the standard error of the mean (SEM). (**B**) Overlaid ^1^H,^15^N TROSY-HSQC spectra of uniformly labeled ^2^H,^13^C,^15^N-labeled β-clamp in the absence (black) and presence (red) of a five-fold molar excess of the QVEMEF peptide. (**C**) Chemical shift perturbations (CSPs) observed upon peptide titration. Asterisks indicate residues with notable CSPs, corresponding to labeled peaks in panel B and mapped structurally in panel D. (**D**) Mapping of affected residues onto the β-clamp structure, bound to the C-terminal Pol III peptide (green) (PDB ID: 3D1F). Residues showing the largest CSPs are labeled, and those that disappear during titration—indicative of intermediate exchange—are shown in blue (Leu177, Tyr244, Val247, Ile317, Ser345, Gln348, Val361, Met364, Arg365). Unassigned residues (including prolines) are labeled grey. DP, differential power; ITC, isothermal titration calorimetry; ^1^H,^15^N TROSY-HSQC, two-dimensional transverse relaxation optimized spectroscopy heteronuclear single quantum coherence experiment correlating ^1^H and ^15^N nuclei.

To confirm whether QVEMEF engages the canonical binding site pocket on β-clamp, we performed two-dimensional NMR studies. We recorded ^1^H,^15^N TROSY-HSQC spectra of ^2^H,^13^C, and ^15^N-labeled β-clamp in the presence and absence of the QVEMEF peptide (Fig. 4B) (See Materials and Methods for details). A titration series was conducted to monitor chemical shift perturbations (CSPs), which revealed widespread peak movements across the β-clamp sequence (Fig. 4C). This pattern, consistent with that observed for other CBMs, suggests conformational changes and altered dynamics upon ligand binding (39). The largest CSPs localized to residues surrounding the canonical CBM binding pocket, including L177, Y244, V247, I317, S345, Q348, V361, M364, and R365. These residues became untraceable during titration due to intermediate exchange on the NMR timescale, indicating their proximity to the binding interface (Fig. 4D).

Together, these data demonstrate that the QVEMEF motif in RecB binds specifically to β-clamp at the canonical CBM binding site with an affinity typical of other known CBMs. Based on these results, we focused on the QVEMEF motif for further functional investigation.

### Mutating the QVEMEF CBM (F1023A) reduces survival after DNA damage

After establishing that the QVEMEF CBM of RecB can bind to β-clamp, we wanted to check the significance of this interaction on the survival of *E. coli* cells after DNA damage. We performed plasmid complementation studies in a *ΔrecB* background using pUT18C constructs harboring either wild type RecB (RecB_WT_) or RecB with an F1023A substitution in QVEMEF (RecB_F1023A_), as this mutation demonstrated loss of binding in NMR studies (Fig. 3D). Expression was induced using IPTG (0.5 mM). The cells were challenged with increasing doses of ciprofloxacin or UV. We found that expressing RecB_WT_ fully restores *ΔrecB* strain survival to the wild-type strain (BTH101) level at all doses of ciprofloxacin (Fig. 5A) or UV (Fig. 5B) tested. In contrast, cells expressing RecB_F1023A_ exhibited a 100-fold reduction in survival following treatment with either 16 ng/mL ciprofloxacin or 30 J/m^2^ UV irradiation (Fig. 5A and B, respectively). It is unlikely that this effect arises from impaired complex formation caused by the mutation, as this would be expected to yield a survival profile comparable to that of the *ΔrecB* strain.

**Figure 5.**
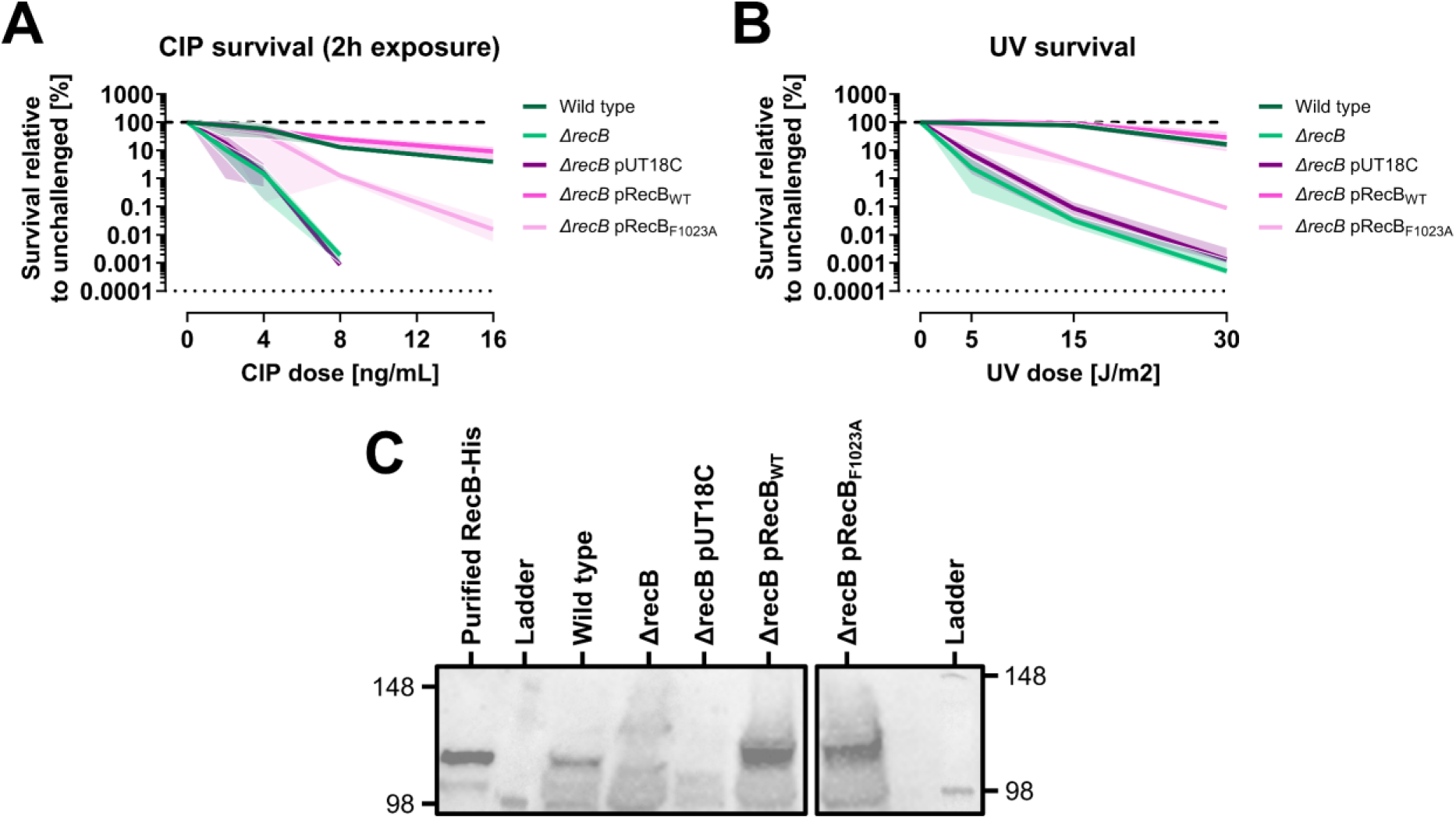
Mutation of the RecB QVEMEF clamp-binding motif impairs survival following DNA damage. (**A** and **B**) Survival of BTH101 *ΔrecB* cells expressing wild-type RecB (RecBWT) or F1023A-mutated RecB (RecBF1023A) from pUT18C plasmids after varying doses of (A) ciprofloxacin (CIP) exposure (0, 2, 4, 8, or 16 ng/mL) for two hours or (B) UV irradiation (0, 5, 15, or 30 J/m^2^). Wild type BTH101, as well as *ΔrecB* strains with or without empty pUT18C vectors, served as controls. Relative survival was calculated through comparison with unchallenged parallels. Solid lines represent means from 2-3 biological replicates; shaded regions indicate standard deviation. Thick dashed lines show survival of unchallenged parallels; thin dotted lines mark the detection limit. (**C**) Western blot analysis of RecB expression from pUT18C constructs in the BTH101 *ΔrecB* background. Each well was loaded with 80 µg protein extract. The two panels shown were cropped from the same gel and processed identically.

To ensure that the decreased survival was not caused by dramatic changes in the protein level of RecB_F1023A_ compared to endogenous RecB or RecB_WT_, we performed Western blotting. We found that the level of RecB_F1023A_ was slightly higher than that of endogenous RecB (wild-type strain), but comparable to that of RecB_WT_ (Fig. 5C). Since both the plasmid-free wild type strain and the strain expressing RecB_WT_ exhibited comparable survival, the reduced survival of the RecB_F1023A_ strain cannot be attributed to protein level differences. In summary, these data indicate a role of the RecB nuclease domain interaction with β-clamp in survival after DNA damage.

### The F1023A CBM mutation impairs the DNA degradation activity of RecBCD

RecBCD plays a dual role in bacterial DNA metabolism (reviewed in (23,54,55)). Upon binding to a double-stranded DNA end, RecBCD initiates processive DNA unwinding and degradation, acting as a potent helicase-nuclease complex. When encountering a Chi site (5′-GCTGGTGG-3′), its activity shifts: degradation is halted, a 3′ overhang is generated, and RecBCD promotes homologous recombination by facilitating RecA loading onto the single-stranded DNA (21-23). This molecular switch—from degradation to recombination—is central to DNA double-strand break repair in *E. coli*. However, in the absence of RecA, this switch cannot occur. As a result, RecBCD continues to degrade DNA unchecked, leading to the production of anucleate cells—cells devoid of chromosomal DNA—following DNA damage (56,57). This RecA-deficient degradation mode is often referred to as “reckless” DNA degradation (58). We leveraged this phenomenon as a functional assay to test whether interaction between RecB and β-clamp influences the reckless degradation by RecBCD. Specifically, we hypothesized that if β-clamp modulates RecBCD activity, then disrupting their interaction might alter the extent of reckless degradation in a *ΔrecA* mutant background.

A *ΔrecA ΔrecB* strain was transformed with either empty plasmid (pUT18C), pRecB_WT_, or pRecB_F1023A_. As a positive control for DNA degradation, a *ΔrecA* single-gene deletion strain was included. All strains were cultured to exponential phase (OD_600_ of 0.2) upon which each culture was split into two aliquots: one aliquot was fixed immediately whereas the second aliquot was exposed to UV irradiation at a dose of 50 J/m^2^. Following UV treatment, cells were allowed to recover for 3 hours prior to fixation in ethanol. DNA was stained using Hoechst 33258 and visualized using fluorescence microscopy.

As expected, the *ΔrecA* cells exhibited extensive DNA degradation, with most cells showing little detectable DNA three hours post-UV (Fig. 6A). Quantitative analysis revealed that *ΔrecA* cells retained only around 16% of the DNA, relative to unchallenged cells (Fig. 6B). The *ΔrecA ΔrecB* double-deletion strain showed a lower reduction in DNA content post-UV (Fig. 6A), retaining about 35%, confirming the role of RecBCD in DNA degradation. Complementation with pRecB_WT_ restored the reckless degradation phenotype, comparable to that of the *ΔrecA* strain (Fig. 6B). Notably, expression of the RecB_F1023A_ mutant reduced DNA degradation compared to wild-type RecB in a highly significant manner (*P*=0.0005) (Fig. 6B). Introduction of the empty plasmid (pUT18C) into the *ΔrecA ΔrecB* strain did not significantly affect DNA content relative to the uncomplemented mutant. In cultures not exposed to UV, the RecB_F1023A_ mutation did not significantly affect the DNA content when compared to wild type RecB expressed from pUT18C (Supplementary Fig. S4). These results may suggest that the interaction between RecB and β-clamp contributes to reckless degradation activity of RecBCD when RecA is absent.

**Figure 6.**
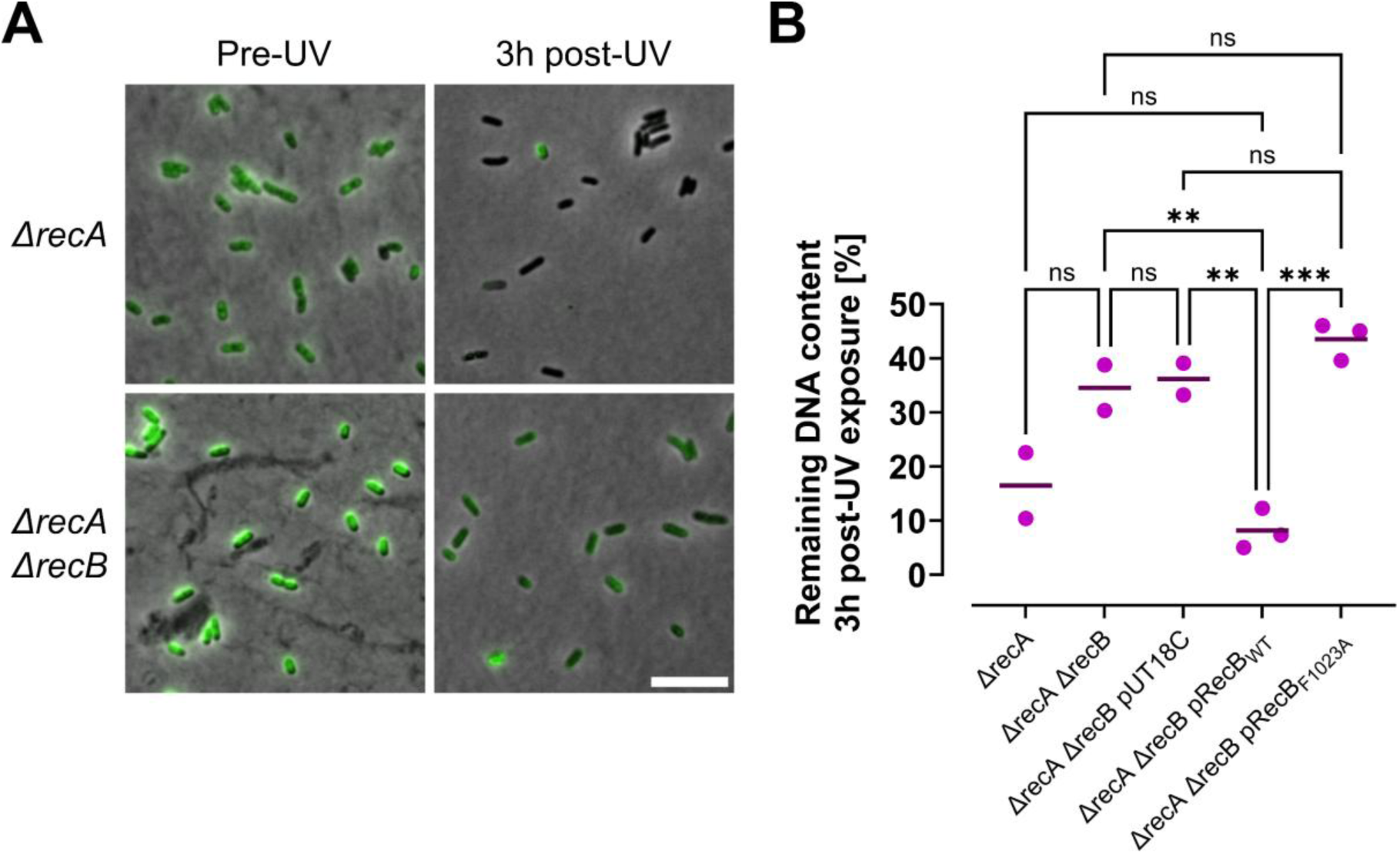
Mutation of the RecB QVEMEF clamp-binding motif (F1023A) impairs DNA degradation following UV irradiation in a BTH101 *ΔrecA* background. (**A**) Representative images of Hoechst 33258-stained (green) *ΔrecA* and *ΔrecA ΔrecB* cells, fixed either before or 3 hours after UV irradiation (50 J/m²). (**B**) Quantification of remaining DNA content post-UV exposure, normalized to unchallenged controls. The *ΔrecA ΔrecB* cells were complemented with either wild-type RecB (pRecBWT) or F1023A-mutated RecB (pRecBF1023A) expressed from pUT18C plasmids. Empty pUT18C plasmids were used for controls. Comparisons were made using ordinary one-way ANOVA with Šidák correction. Lines represent normalized medians from 2-3 biological replicates; dots indicate individual replicates. ns, non-significant; \*\**P* ≤ 0.01; \*\*\**P* ≤ 0.001.

## DISCUSSION

In this study, we identified and characterized a novel interaction between the *E. coli* β-clamp and the RecBCD complex. Using a combination of bacterial two-hybrid assays, co-immunoprecipitation, fluorescence microscopy, and NMR spectroscopy, we discovered that the interaction involves a CBM in the nuclease domain of RecB (QVEMEF). We also found the interaction of β-clamp with RecBCD to be functionally relevant for RecBCD’s degradation activity and contribute to cell survival following DNA damage. These findings uncover a previously unrecognized link between β-clamp and DSB repair in *E. coli*, as well as possibly RecBCD-mediated salvage of stalled or collapsed replication forks.

### RecBCD associates with β-clamp during unchallenged growth

Using fluorescence microscopy, we showed that around 32% of unchallenged cells displayed RecB and β-clamp foci within close proximity of one another, indicating spatial colocalization under normal growth conditions (Fig. 1C and D). This observation aligns with the view that RecBCD is not exclusively a damage-induced responder but also plays constitutive roles in safeguarding DNA replication fidelity (46).

In unchallenged conditions, β-clamp is largely associated with active replication forks, where it functions as a processivity factor and as a hub for recruiting various enzymes for DNA metabolism. The colocalization of RecBCD with β-clamp under these conditions raises the possibility that RecBCD may localize near, or be recruited to, replication forks prior to overt DNA damage. Given that replication forks are frequently challenged by endogenous obstacles—such as DNA secondary structures, tightly bound proteins, or template lesions (47)—it is plausible that RecBCD is recruited in a preemptive or surveillance capacity to help resolve impediments to fork progression. Alternatively, the observed level of colocalization may reflect replication forks actively undergoing restart or repair. This interpretation is supported by prior evidence in *E. coli* that replication fork stalling occurs multiple times per replication cycle and can lead to fork regression or spontaneous DSB formation (48,49). In these contexts, RecBCD is known to engage either RecA-dependent homologous recombination or RecA-independent pathways to facilitate fork restart (46).

Our data indicate that the RecB–β-clamp interaction persists even in a *ΔrecA* background (Supplementary Fig. S1), albeit with delayed detection, suggesting that the interaction does not strictly require the homologous recombination machinery. Instead, it may occur upstream of recombination or reflect a broader role for RecBCD in monitoring or responding to replication stress.

### Implications of RecBCD–β-clamp interactions in response to DNA damage

We also observed colocalization between RecB and β-clamp following ciprofloxacin treatment (Fig. 1D). Although the fraction of colocalizing cells was slightly reduced after treatment, this decrease coincided with a notable reduction in the average number of β-clamp foci per cell (Supplementary Fig. S2B)—likely reflecting widespread replisome disassembly upon drug exposure (50). Under these conditions, persistent β-clamp foci likely represent β-clamps retained on DNA after replisome collapse, either at sites of unresolved forks or behind previously active replication forks.

Ciprofloxacin at the minimum inhibitory concentration (20 ng/mL) affects the survival of cells in a time-dependent manner through induction of substantial DNA damage (37). The DNA damage includes replication-dependent and replication-independent DSBs (59,60), both of which necessitate repair through homologous recombination pathways involving RecBCD and RecA. Although our microscopy data cannot determine whether the RecB–β-clamp interaction occurs at replication forks during DSB repair, at distal damage sites, or primarily under normal growth conditions, additional evidence suggests its role extends beyond DNA replication. First, after UV or ciprofloxacin challenge, we demonstrated 100-fold lowered survival of a strain expressing RecB deficient in β-clamp binding (Fig. 5). Second, we performed an experiment using the BACTH assay, which indicated an increased RecB–β-clamp interaction following UV irradiation (15 J/m²) (Supplementary Fig. S5). Together, these findings support a model in which RecBCD–β-clamp interactions are not restricted to unperturbed replication forks but also occur in DNA repair contexts, potentially involving β-clamp that remains bound to DNA after replisome disassembly.

### Multiple Candidate Clamp-Binding Motifs in RecB

We initially identified two candidate CBMs within RecB: QALRF (helicase domain) and QVEMEF (nuclease domain). Only the latter was validated to interact with β-clamp by NMR spectroscopy (Fig. 3), though both motifs appeared to support β-clamp interaction in the BACTH system when tested in truncated RecB constructs (Fig. 2C).

It may be that complete alanine substitution of QALRF disrupts proper helicase domain folding or prevents RecBCD complex assembly, which indirectly leads to loss of interaction. As demonstrated, RecC and RecD subunits are required for detectable interaction in the BACTH assay (Fig. 1A). It could also be that the QALRF motif may require specific conformational states, post-translational modifications, or cofactor binding of RecBCD for β-clamp recognition—conditions that are not replicated in the NMR experimental setup. Alternatively, the binding affinity was outside the range detectable for STD-NMR (around 10^-8^ M to 10^-3^ M). Although QALRF is a weaker match to the canonical CBM consensus sequence compared to QVEMEF (1), its similarity to the internal CBM of Pol III (QADMF) suggests possible evolutionary or structural relevance. Given that Pol III possesses both internal and C-terminal CBMs (2,12,53) it is plausible that RecB also employs a dual-motif architecture or that other regions not identified here participate and stabilize the interaction. Indeed, multiple binding surfaces have been described on β-clamp, allowing for simultaneous or sequential interaction with various client proteins (61-63). It is therefore conceivable that RecB, either alone or in the context of the RecBCD complex, engages the clamp via additional, noncanonical interfaces or transient contacts that escape detection by motif searches or in vitro assays. Further investigation, including structural and mutational analyses beyond canonical CBM sequences, will be needed to fully map the interaction interface.

The QVEMEF motif aligns well with key features of known CBMs, including a conserved glutamine at position 1, an aliphatic residue at position 2, a methionine at position 4, and an essential phenylalanine at the last position. Its validation by NMR and its functional importance in β-clamp binding and nuclease activity underscore its biological relevance.

### Conformational regulation of RecBCD and the role of the QVEMEF motif

The QVEMEF motif resides within the nuclease domain of RecB, a region buried in the crystal structure of the DNA-bound RecBCD complex (Fig. 2B). This positioning suggests that β-clamp interaction via this motif requires structural rearrangement. According to the swing model (64), Chi recognition triggers a dramatic conformational shift in RecBCD: the RecB nuclease domain swings outward on its 19 amino-acid linker—moving from its initial tucked position to emerge near the RecC tunnel exit—thereby potentially exposing motifs such as that involved in β-clamp binding and enabling the switch from DNA degradation to recombination initiation.

Recent structural and biochemical evidence complicates this model. Pavankumar et al. (65) demonstrated that the nuclease domain remains in its pre-Chi conformation until the 5′ DNA tail exceeds approximately 10 nucleotides. Only then do more extensive conformational shifts occur—likely involving partial loosening rather than full domain rotation. These intermediate states appear sufficient to support RecA loading and modulate enzymatic activity without full domain detachment.

In this context, exposure of the QVEMEF motif may be dynamically regulated by DNA substrate structure and Chi recognition. Such conditional accessibility could modulate β-clamp binding, potentially stabilizing a recombination-competent conformation or tuning RecBCD activity in response to replication stress or specific DNA intermediates. Current structural models of RecBCD are based on its DNA-bound conformation, where the RecB nuclease domain is buried and the QVEMEF motif is inaccessible. However, β-clamp may bind RecBCD in a DNA-free or partially unbound state—such as at free DNA ends or prior to re-engagement with a substrate—where alternative conformations expose interaction motifs. Since no structures of the full, unbound RecBCD complex exist, it is plausible that β-clamp binding reflects a conformation distinct from those captured crystallographically, possibly inducing or stabilizing a recombination-ready state.

Our BACTH data support direct interaction between the QVEMEF motif and β-clamp (Fig. 2C), as mutation of the motif abolishes the interaction. NMR analysis further confirms this interaction and shows that a single F1023A substitution is sufficient to disrupt binding (Fig. 3C and D). Several residues in QVEMEF—particularly valine and the two glutamates— have been implicated in RecA binding (24), but the phenylalanine has not. Additionally, a RecB mutation (D1080A), which disrupts the interaction between RecB and RecA, has been demonstrated to lead to survival rates after UV exposure that are comparable to those observed in strains with a complete *recB* deletion (66). Thus, the observed, moderate survival defect (Fig. 5) is unlikely to result from impaired RecA interaction.

Structural models suggest that the phenylalanine may also contact RecC in the pre-Chi conformation (19). However, like the case of RecA interaction, complete disruption of the RecB–RecC contact would likely also phenocopy a *ΔrecB* strain, which shows a much more severe survival defect than our mutant (Fig. 5A and B). The moderate (∼100-fold) reduction in survival we observe, along with impaired DNA degradation (Fig. 6), may align more closely with the loss of β-clamp binding. If this interpretation is accurate, it raises an intriguing possibility: that β-clamp binding after Chi recognition enhances RecBCD’s processivity, potentially accounting for the extensive DNA degradation observed in *ΔrecA* mutants.

Given that the QVEMEF motif mediates interactions with both RecA and β-clamp, these partners may bind in a mutually exclusive manner. This could reflect conformational or temporal switching, allowing RecBCD to dynamically coordinate between DNA degradation and homologous recombination during the DNA damage response. Alternatively, these interactions may operate in a coordinated or sequential manner. For example, clamp binding might help stabilize RecBCD at DNA ends or stalled forks, positioning the complex for a timely handoff to RecA. In this scenario, β-clamp and RecA could act in concert to facilitate an efficient transition from DNA processing to strand invasion—supporting RecA’s homology search and filament formation on the processed 3′ ssDNA tail.

Our study identifies a previously unrecognized functional interaction between β-clamp and the RecBCD complex, mediated by a novel CBM (QVEMEF) in RecB. This interaction appears conformationally regulated and may modulate the recruitment of RecBCD and subsequent nuclease activity at DNA breaks. Beyond its canonical role in replication, β-clamp may act as a scaffold for coordinating repair processes—stabilizing RecBCD at stalled forks or persistent double-strand breaks and ensuring spatial control during recombination. These findings further illuminate the multifunctionality of the sliding clamp, as shown for PCNA, expanding its role in coordinating replication and repair pathways during the bacterial DNA damage response. Disrupting the RecB–β-clamp interaction could present novel targets for antimicrobial intervention. Further work is needed to resolve the structural details and assess the physiological relevance of the interaction of β-clamp with RecBCD.

## DATA AVAILABILITY

Data underlying this article will be shared on reasonable request to the corresponding author.

## SUPPLEMENTARY DATA

Supplementary data is available online at bioRxiv.

## AUTHOR CONTRIBUTIONS

Conceptualization: I.M.M.R., S.B.R., K.S., E.H.; Methodology: I.M.M.R., S.B.R., S.S., K.V., P.H.B., L.J., M.B., J.A.B., B.K., K.S., E.H.; Investigation: I.M.M.R. S.B.R., S.S., K.V.; Formal analysis: I.M.M.R., S.B.R., S.S., K.V.; Visualization: I.M.M.R., S.B.R., S.S., K.V., P.H.B.; Funding acquisition: M.B., J.A.B., B.B.K., K.S., E.H.; Supervision: M.B., J.A.B., B.B.K., K.S., E.H.; Writing — original draft: I.M.M.R., S.B.R., E.H.; Writing — review & editing: all authors.

## ACKNOWLEDGEMENTS

We are grateful to Dr. Meriem El Karoui (University of Edinburgh) for providing the RecB-HaloTag construct and advising on experimental setup for imaging of RecB. We thank Prof. Tsutomu Katayama (University of Kyushu) for providing GFP-RecN, and Prof. Michael T. Laub (MIT and Howard Hughes Medical Institute) for the pBAD30 plasmid. We thank previous group members at OUH Anne Wahl for assistance with strain construction, and Christiane Nørgaard for BACTH method optimization during initial phases of the project. We thank Signe A. Sjørup (University of Copenhagen) for technical support and Andreas Prestel (University of Copenhagen) for NMR advice. The graphical abstract was created in BioRender (https://BioRender.com/9puqdqj). We acknowledge the assistance of ChatGPT by OpenAI in providing suggestions on phrasing. The human authors performed all editing.

## REFERENCES

1. Simonsen, S., Søgaard, C.K., Olsen, J.G., Otterlei, M. and Kragelund, B.B. (2024) The bacterial DNA sliding clamp, β-clamp: structure, interactions, dynamics and drug discovery. Cell Mol Life Sci, 81, 245.

2. López de Saro, F.J. and O’Donnell, M. (2001) Interaction of the beta sliding clamp with MutS, ligase, and DNA polymerase I. Proc Natl Acad Sci U S A, 98, 8376–8380.

3. Stukenberg, P.T., Studwell-Vaughan, P.S. and O’Donnell, M. (1991) Mechanism of the sliding beta-clamp of DNA polymerase III holoenzyme. J Biol Chem, 266, 11328–11334.

4. Katayama, T., Kubota, T., Kurokawa, K., Crooke, E. and Sekimizu, K. (1998) The initiator function of DnaA protein is negatively regulated by the sliding clamp of the E. coli chromosomal replicase. Cell, 94, 61–71.

5. Kurz, M., Dalrymple, B., Wijffels, G. and Kongsuwan, K. (2004) Interaction of the sliding clamp beta-subunit and Hda, a DnaA-related protein. J Bacteriol, 186, 3508–3515.

6. Hughes, A.J., Jr., Bryan, S.K., Chen, H., Moses, R.E. and McHenry, C.S. (1991) Escherichia coli DNA polymerase II is stimulated by DNA polymerase III holoenzyme auxiliary subunits. J Biol Chem, 266, 4568–4573.

7. Lenne-Samuel, N., Wagner, J., Etienne, H. and Fuchs, R.P. (2002) The processivity factor beta controls DNA polymerase IV traffic during spontaneous mutagenesis and translesion synthesis in vivo. EMBO Rep, 3, 45–49.

8. Tang, M., Shen, X., Frank, E.G., O’Donnell, M., Woodgate, R. and Goodman, M.F. (1999) UmuD’(2)C is an error-prone DNA polymerase, Escherichia coli pol V. Proc Natl Acad Sci U S A, 96, 8919–8924.

9. Wagner, J., Fujii, S., Gruz, P., Nohmi, T. and Fuchs, R.P. (2000) The beta clamp targets DNA polymerase IV to DNA and strongly increases its processivity. EMBO Rep, 1, 484–488.

10. Henry, C., Kaur, G., Cherry, M.E., Henrikus, S.S., Bonde, N.J., Sharma, N., Beyer, H.A., Wood, E.A., Chitteni-Pattu, S., van Oijen, A.M., et al. (2023) RecF protein targeting to post-replication (daughter strand) gaps II: RecF interaction with replisomes. Nucleic Acids Res, 51, 5714–5742.

11. Indiani, C., McInerney, P., Georgescu, R., Goodman, M.F. and O’Donnell, M. (2005) A sliding-clamp toolbelt binds high- and low-fidelity DNA polymerases simultaneously. Mol Cell, 19, 805–815.

12. Dalrymple, B.P., Kongsuwan, K., Wijffels, G., Dixon, N.E. and Jennings, P.A. (2001) A universal protein-protein interaction motif in the eubacterial DNA replication and repair systems. Proc Natl Acad Sci U S A, 98, 11627–11632.

13. Altieri, A.S. and Kelman, Z. (2018) DNA Sliding Clamps as Therapeutic Targets. Front Mol Biosci, 5, 87.

14. Olaisen, C., Kvitvang, H.F.N., Lee, S., Almaas, E., Bruheim, P., Drabløs, F. and Otterlei, M. (2018) The role of PCNA as a scaffold protein in cellular signaling is functionally conserved between yeast and humans. FEBS Open Bio, 8, 1135–1145.

15. Prestel, A., Wichmann, N., Martins, J.M., Marabini, R., Kassem, N., Broendum, S.S., Otterlei, M., Nielsen, O., Willemoës, M., Ploug, M. et al. (2019) The PCNA interaction motifs revisited: thinking outside the PIP-box. Cell Mol Life Sci, 76, 4923–4943.

16. Cardano, M., Tribioli, C. and Prosperi, E. (2020) Targeting Proliferating Cell Nuclear Antigen (PCNA) as an Effective Strategy to Inhibit Tumor Cell Proliferation. Curr Cancer Drug Targets, 20, 240–252.

17. Li, J., Holzschu, D.L. and Sugiyama, T. (2013) PCNA is efficiently loaded on the DNA recombination intermediate to modulate polymerase δ, η, and ζ activities. Proc Natl Acad Sci U S A, 110, 7672–7677.

18. Li, X., Stith, C.M., Burgers, P.M. and Heyer, W.D. (2009) PCNA is required for initiation of recombination-associated DNA synthesis by DNA polymerase delta. Mol Cell, 36, 704–713.

19. Amundsen, S.K. and Smith, G.R. (2023) RecBCD enzyme: mechanistic insights from mutants of a complex helicase-nuclease. Microbiol Mol Biol Rev, 87, e0004123.

20. Kowalczykowski, S.C. (2000) Initiation of genetic recombination and recombination-dependent replication. Trends Biochem Sci, 25, 156–165.

21. Anderson, D.G. and Kowalczykowski, S.C. (1997) The recombination hot spot chi is a regulatory element that switches the polarity of DNA degradation by the RecBCD enzyme. Genes Dev, 11, 571–581.

22. Dixon, D.A. and Kowalczykowski, S.C. (1993) The recombination hotspot chi is a regulatory sequence that acts by attenuating the nuclease activity of the E. coli RecBCD enzyme. Cell, 73, 87–96.

23. Smith, G.R. (2012) How RecBCD enzyme and Chi promote DNA break repair and recombination: a molecular biologist’s view. Microbiol Mol Biol Rev, 76, 217–228.

24. Spies, M. and Kowalczykowski, S.C. (2006) The RecA binding locus of RecBCD is a general domain for recruitment of DNA strand exchange proteins. Mol Cell, 21, 573–580.

25. Battesti, A. and Bouveret, E. (2012) The bacterial two-hybrid system based on adenylate cyclase reconstitution in Escherichia coli. Methods, 58, 325–334.

26. Mehla, J., Caufield, J.H., Sakhawalkar, N. and Uetz, P. (2017) In Shukla, A. K. (ed.), Methods in Enzymology. Academic Press, Vol. 586, pp. 333-358.

27. Thomason, L.C., Costantino, N. and Court, D.L. (2007) E. coli genome manipulation by P1 transduction. Curr Protoc Mol Biol, 79, 1.17.01-08.

28. Baba, T., Ara, T., Hasegawa, M., Takai, Y., Okumura, Y., Baba, M., Datsenko, K.A., Tomita, M., Wanner, B.L. and Mori, H. (2006) Construction of Escherichia coli K-12 in-frame, single-gene knockout mutants: the Keio collection. Mol Syst Biol, 2, 2006.0008.

29. Cherepanov, P.P. and Wackernagel, W. (1995) Gene disruption in Escherichia coli: TcR and KmR cassettes with the option of Flp-catalyzed excision of the antibiotic-resistance determinant. Gene, 158, 9–14.

30. Karimova, G., Pidoux, J., Ullmann, A. and Ladant, D. (1998) A bacterial two-hybrid system based on a reconstituted signal transduction pathway. Proceedings of the National Academy of Sciences, 95, 5752–5756.

31. Biek, D.P. and Cohen, S.N. (1986) Identification and characterization of recD, a gene affecting plasmid maintenance and recombination in Escherichia coli. Journal of Bacteriology, 167, 594–603.

32. Lepore, A., Taylor, H., Landgraf, D., Okumus, B., Jaramillo-Riveri, S., McLaren, L., Bakshi, S., Paulsson, J. and Karoui, M.E. (2019) Quantification of very low-abundant proteins in bacteria using the HaloTag and epi-fluorescence microscopy. Scientific Reports, 9, 7902.

33. Ozaki, S., Matsuda, Y., Keyamura, K., Kawakami, H., Noguchi, Y., Kasho, K., Nagata, K., Masuda, T., Sakiyama, Y. and Katayama, T. (2013) A replicase clamp-binding dynamin-like protein promotes colocalization of nascent DNA strands and equipartitioning of chromosomes in E. coli. Cell Rep, 4, 985–995.

34. Guzman, L.M., Belin, D., Carson, M.J. and Beckwith, J. (1995) Tight regulation, modulation, and high-level expression by vectors containing the arabinose PBAD promoter. J Bacteriol, 177, 4121–4130.

35. Johnsen, L., Flåtten, I., Morigen, Dalhus, B., Bjørås, M., Waldminghaus, T. and Skarstad, K. (2011) The G157C mutation in the Escherichia coli sliding clamp specifically affects initiation of replication. Mol Microbiol, 79, 433–446.

36. Nedal, A., Ræder, S.B., Dalhus, B., Helgesen, E., Forstrøm, R.J., Lindland, K., Sumabe, B.K., Martinsen, J.H., Kragelund, B.B., Skarstad, K. et al. (2020) Peptides containing the PCNA interacting motif APIM bind to the β-clamp and inhibit bacterial growth and mutagenesis. Nucleic Acids Res, 48, 5540–5554.

37. Vikedal, K., Ræder, S.B., Riisnæs, I.Mathilde M., Bjørås, M., Booth, J.A., Skarstad, K. and Helgesen, E. (2025) RecN and RecA orchestrate an ordered DNA supercompaction response following ciprofloxacin-induced DNA damage in Escherichia coli. Nucleic Acids Research, 53, gkaf437.

38. Ducret, A., Quardokus, E.M. and Brun, Y.V. (2016) MicrobeJ, a tool for high throughput bacterial cell detection and quantitative analysis. Nat Microbiol, 1, 16077.

39. Simonsen, S., Prestel, A., Østerlund, E.C., Otterlei, M., Jørgensen, T.J.D. and Kragelund, B.B. (2025) Responses to Ligand Binding in the Bacterial DNA Sliding Clamp "β-Clamp" Manifest in Dynamic Allosteric Effects. Proteins.

40. Gasteiger E., H.C., Gattiker A., Duvaud S., Wilkins M. R., Appel R. D., Bairoch A. (2005) In Walker, J. M. (ed.). Humana Press.

41. Vranken, W.F., Boucher, W., Stevens, T.J., Fogh, R.H., Pajon, A., Llinas, M., Ulrich, E.L., Markley, J.L., Ionides, J. and Laue, E.D. (2005) The CCPN data model for NMR spectroscopy: development of a software pipeline. Proteins, 59, 687–696.

42. Mulder, F.A., Schipper, D., Bott, R. and Boelens, R. (1999) Altered flexibility in the substrate-binding site of related native and engineered high-alkaline Bacillus subtilisins. J Mol Biol, 292, 111–123.

43. Schindelin, J., Arganda-Carreras, I., Frise, E., Kaynig, V., Longair, M., Pietzsch, T., Preibisch, S., Rueden, C., Saalfeld, S., Schmid, B. et al. (2012) Fiji: an open-source platform for biological-image analysis. Nature Methods, 9, 676-682.

44. Lepore, A., Thédié, D., McLaren, L., Goossens, L., Azeroglu, B., Pambos, O.J., Kapanidis, A.N. and El Karoui, M. (2025) In vivo single-molecule imaging of RecB reveals efficient repair of DNA damage in Escherichia coli. Nucleic Acids Research, 53, gkaf454.

45. 45. Moolman, M.C., Krishnan, S.T., Kerssemakers, J.W.J., van den Berg, A., Tulinski, P., Depken, M., Reyes-Lamothe, R., Sherratt, D.J. and Dekker, N.H. (2014) Slow unloading leads to DNA-bound β2-sliding clamp accumulation in live Escherichia coli cells. Nature Communications, 5, 5820.

46. Michel, B., Sinha, A.K. and Leach, D.R.F. (2018) Replication Fork Breakage and Restart in Escherichia coli. Microbiol Mol Biol Rev, 82, e00013–00018.

47. Mirkin, E.V. and Mirkin, S.M. (2007) Replication fork stalling at natural impediments. Microbiol Mol Biol Rev, 71, 13–35.

48. Michel, B. and Sandler, S.J. (2017) Replication Restart in Bacteria. J Bacteriol, 199, e00102–00117.

49. Sinha, A.K., Possoz, C., Durand, A., Desfontaines, J.-M., Barre, F.-X., Leach, D.R.F. and Michel, B. (2018) Broken replication forks trigger heritable DNA breaks in the terminus of a circular chromosome. PLOS Genetics, 14, e1007256.

50. Ojkic, N., Lilja, E., Direito, S., Dawson, A., Allen, R.J. and Waclaw, B. (2020) A Roadblock-and-Kill Mechanism of Action Model for the DNA-Targeting Antibiotic Ciprofloxacin. Antimicrob Agents Chemother, 64, e02487–02419.

51. Kalita, I., Iosub, I.A., McLaren, L., Goossens, L., Granneman, S. and El Karoui, M. (2025) An Hfq-dependent post-transcriptional mechanism fine tunes RecB expression in Escherichia coli. eLife, 13, RP94918.

52. Payne-Dwyer, A.L., Syeda, A.H., Shepherd, J.W., Frame, L. and Leake, M.C. (2022) RecA and RecB: probing complexes of DNA repair proteins with mitomycin C in live Escherichia coli with single-molecule sensitivity. J R Soc Interface, 19, 20220437.

53. Dohrmann, P.R. and McHenry, C.S. (2005) A bipartite polymerase-processivity factor interaction: only the internal beta binding site of the alpha subunit is required for processive replication by the DNA polymerase III holoenzyme. J Mol Biol, 350, 228–239.

54. Dillingham, M.S. and Kowalczykowski, S.C. (2008) RecBCD enzyme and the repair of double-stranded DNA breaks. Microbiol Mol Biol Rev, 72, 642–671, Table of Contents.

55. Michel, B. and Leach, D. (2012) Homologous Recombination-Enzymes and Pathways. EcoSal Plus, 5, 1–46.

56. Repar, J., Briški, N., Buljubašić, M., Zahradka, K. and Zahradka, D. (2013) Exonuclease VII is involved in "reckless" DNA degradation in UV-irradiated Escherichia coli. Mutat Res, 750, 96–104.

57. Skarstad, K. and Boye, E. (1993) Degradation of individual chromosomes in recA mutants of Escherichia coli. J Bacteriol, 175, 5505–5509.

58. Willetts, N.S. and Clark, A.J. (1969) Characteristics of some multiply recombination-deficient strains of Escherichia coli. J Bacteriol, 100, 231–239.

59. Wentzell, L.M. and Maxwell, A. (2000) The complex of DNA gyrase and quinolone drugs on DNA forms a barrier to the T7 DNA polymerase replication complex. J Mol Biol, 304, 779–791.

60. Zhao, X., Malik, M., Chan, N., Drlica-Wagner, A., Wang, J.Y., Li, X. and Drlica, K. (2006) Lethal action of quinolones against a temperature-sensitive dnaB replication mutant of Escherichia coli. Antimicrob Agents Chemother, 50, 362–364.

61. Chang, S., Laureti, L., Thrall, E.S., Kay, M.S., Philippin, G., Jergic, S., Pagès, V. and Loparo, J.J. (2025) A bipartite interaction with the processivity clamp potentiates Pol IV-mediated TLS. Proceedings of the National Academy of Sciences, 122, e2421471122.

62. Heltzel, J.M., Maul, R.W., Scouten Ponticelli, S.K. and Sutton, M.D. (2009) A model for DNA polymerase switching involving a single cleft and the rim of the sliding clamp. Proc Natl Acad Sci U S A, 106, 12664–12669.

63. Kath, J.E., Jergic, S., Heltzel, J.M., Jacob, D.T., Dixon, N.E., Sutton, M.D., Walker, G.C. and Loparo, J.J. (2014) Polymerase exchange on single DNA molecules reveals processivity clamp control of translesion synthesis. Proc Natl Acad Sci U S A, 111, 7647–7652.

64. Taylor, A.F., Amundsen, S.K., Guttman, M., Lee, K.K., Luo, J., Ranish, J. and Smith, G.R. (2014) Control of RecBCD enzyme activity by DNA binding- and Chi hotspot-dependent conformational changes. J Mol Biol, 426, 3479–3499.

65. Pavankumar, T.L., Wong, C.J., Wong, Y.K., Spies, M. and Kowalczykowski, S.C. (2024) Trans-complementation by the RecB nuclease domain of RecBCD enzyme reveals new insight into RecA loading upon χ recognition. Nucleic Acids Res, 52, 2578–2589.

66. Amundsen, S.K., Taylor, A.F. and Smith, G.R. (2000) The RecD subunit of the Escherichia coli RecBCD enzyme inhibits RecA loading, homologous recombination, and DNA repair. Proc Natl Acad Sci U S A, 97, 7399–7404.

