## Supplementary for "The RecBCD complex interacts directly with the DNA sliding clamp in *Escherichia coli*"

**Table S1.** Hit *E. coli* genes from the yeast two-hybrid screening, classified according to the confidence of their interaction with the β-clamp (*dnaN*). The cellular localization of the gene products is listed. The *recB* hit is highlighted.

| Gene | Confidence in interaction | Cellular localization |
|---|---|---|
| *amiC* | Moderate | Cytoplasm |
| *b0942* | Moderate | Membrane |
| *b1504* | High | Membrane |
| *b1978* | Very high | Membrane |
| *b2074* | Very high | Membrane |
| *b2075* | Good | Membrane |
| *b2225* | Very high | Membrane |
| *b2520* | Moderate | Membrane |
| *b2973* | High | Membrane |
| *bglX* | Very high | Membrane |
| *cysI* | High | Cytoplasm |
| *dnaX* | Moderate | Cytoplasm |
| *eno* | Very high | Membrane |
| *fecA* | Good | Membrane |
| *ftsK* | High | Cytoplasm |
| *fusA* | High | Cytoplasm |
| *infA* | Good | Membrane |

| Gene | Confidence in interaction | Cellular localization |
|---|---|---|
| *ligA* | High | Cytoplasm |
| *lipB* | Moderate | Cytoplasm |
| *mepA* | Moderate | Membrane |
| *modA* | Moderate | Membrane |
| *nmpC* | Very high | Membrane |
| *ompC* | Very high | Membrane |
| *ompN* | Very high | Membrane |
| *pepA* | High | Cytoplasm |
| *pepT* | Good | Cytoplasm |
| *pflA* | Good | Cytoplasm |
| *phoE* | Very high | Membrane |
| *ppsA* | High | Cytoplasm |
| *prpD* | Moderate | Cytoplasm |
| ***recB*** | **Moderate** | **Cytoplasm** |
| *rpe* | Good | Cytoplasm |
| *secD* | Moderate | Membrane |
| *speB* | Very high | Cytoplasm |
| *sufI* | Good | Membrane |
| *talc* | High | Cytoplasm |
| *tdh* | Moderate | Cytoplasm |
| *tolA* | High | Membrane |
| *torC* | High | Membrane |
| *yadM* | Good | Membrane |
| *yagX* | Moderate | Membrane |
| *ydbA_1* | Good | Membrane |
| *yehB* | Moderate | Membrane |
| *ytfN* | Very high | Membrane |
| *yhjN* | Very high | Membrane |
| *yihF* | Moderate | Membrane |
| *yjjQ* | Good | Cytoplasm |

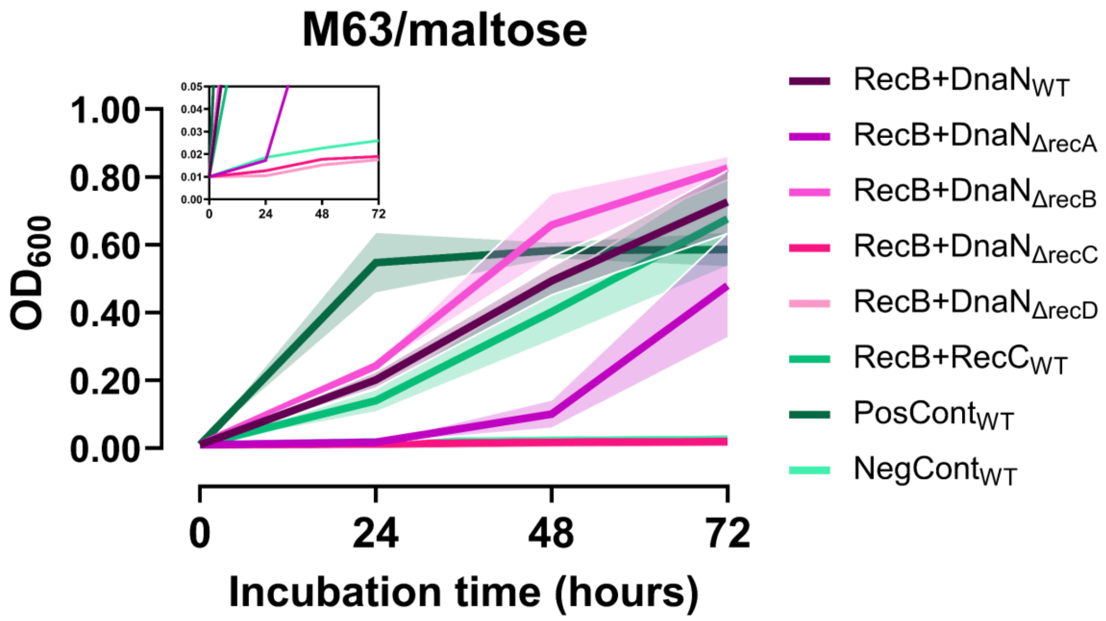


**Figure S1.** BACTH-based interaction between RecB and β-clamp in wild-type and RecBCD mutant backgrounds. Growth of *E. coli* BTH101 strains co-expressing RecB–T18 and β-clamp (DnaN)–T25 fusion proteins was monitored in M63/maltose minimal medium over 72 hours at 30 °C. OD600 was measured at 24, 48, and 72 hours to assess interaction-dependent growth based on reconstitution of adenylate cyclase activity. The RecB–β-clamp interaction was assessed in wild-type BTH101 and in isogenic deletion strains lacking *recA*, *recB*, *recC*, or *recD*. Wild-type cells co-expressing RecB and RecC were included as an internal control for protein–protein interaction. Co-transformed *zip* fusions served as a positive control, and empty vectors (T18 and T25 alone) as a negative control. While interaction-dependent growth was observed in wild-type, *ΔrecA*, and *ΔrecB* strains, the interaction was strongly impaired in both *ΔrecC* and *ΔrecD* backgrounds—suggesting a requirement for intact RecBCD architecture. Lines represent means from three biological replicates; shaded regions indicate SEM.

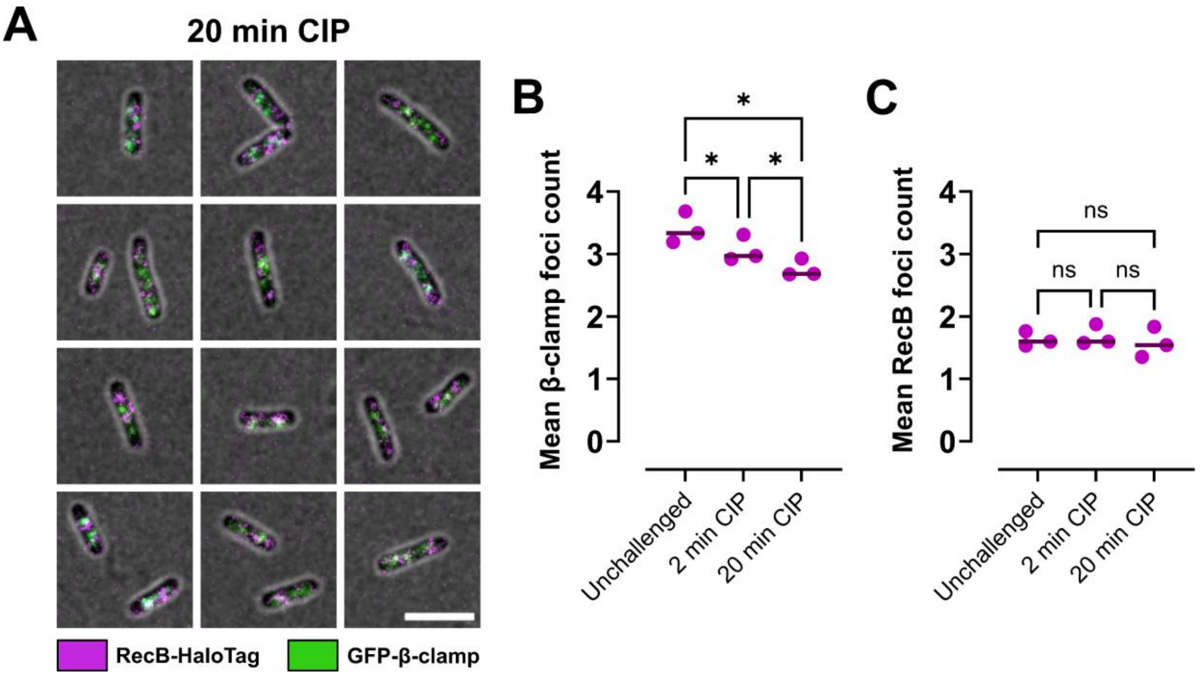


**Figure S2.** Ciprofloxacin treatment reduces β-clamp foci but not RecB foci in live *E. coli* cells. (**A**) Representative fluorescence microscopy images of fixed EH219 cells treated with ciprofloxacin (20 ng/mL, 20 min), showing JF549-labeled RecB–HaloTag (magenta) and endogenous GFP–β-clamp (green). Colocalized signals appear white. (**B**) Quantification of mean β-clamp foci per cell in untreated cells and after 2 or 20 min of ciprofloxacin treatment. (**C**) Quantification of mean RecB foci per cell under the same conditions. Statistical comparisons were made using repeated measures one-way ANOVA with Tukey's multiple comparisons test. Lines represent the mean of three biological replicates; dots represent individual replicate means. ns, not significant; **P* ≤ 0.05.

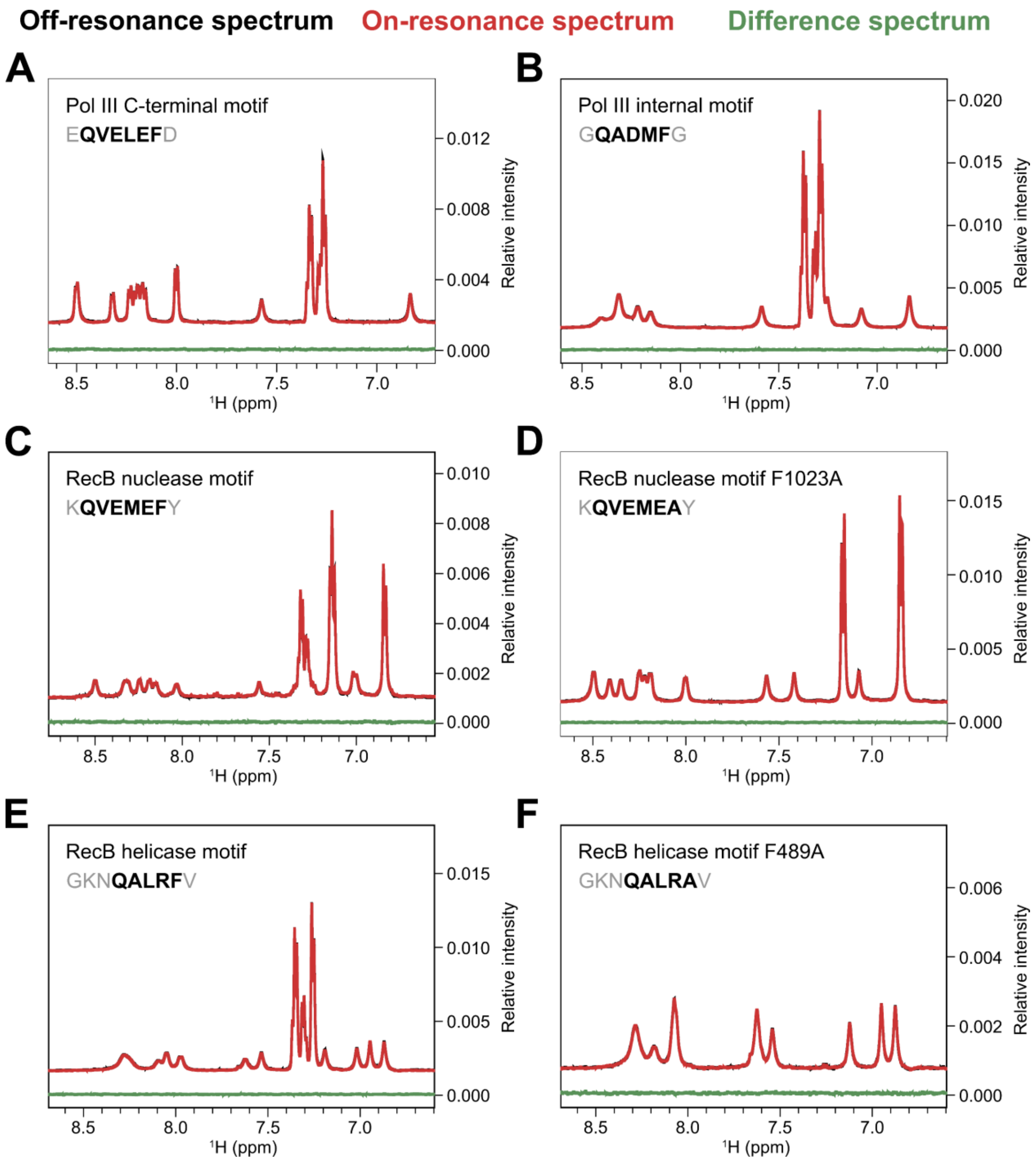


**Figure S3.** Control STD-NMR spectra confirm that peptides do not experience saturation in the absence of β-clamp. (**A-F**) Saturation Transfer Difference (STD) NMR spectra of synthetic peptides in the absence of purified β-clamp protein. For each sample, the off-resonance (reference) spectrum is shown in black, the on-resonance (saturated) spectrum in red, and the resulting STD difference spectrum in green. Peptides tested: (A) EQVELEFD, (B) GQADMFG, (C) KQVEMEFY, (D) KQVEMEAY, (E) GKNQALRFV, and (F) GKNQALRAV. No detectable signals were observed in the difference spectra under the conditions used, confirming that the peptides do not experience saturation in the absence of β-clamp binding.

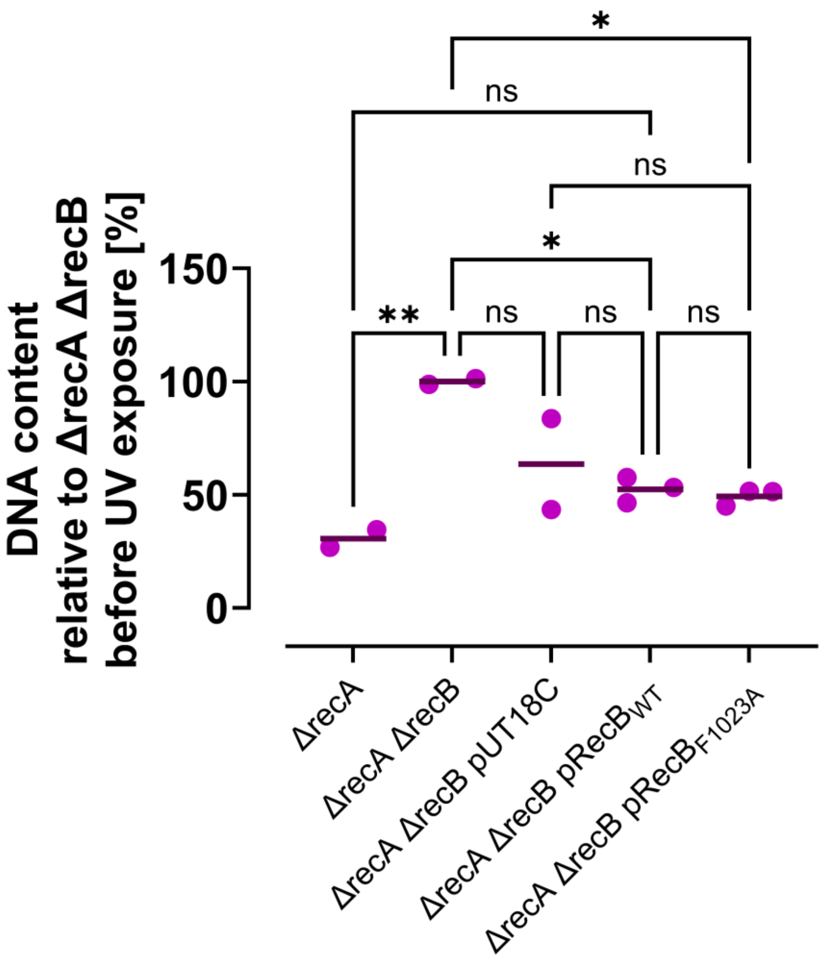


**Figure S4.** The $RecB_{F1023A}$ mutation does not alter DNA content under non-damaging conditions. Cellular DNA content (Y-axis) is shown relative to *ΔrecA ΔrecB* cells prior to UV exposure (%), as measured in strains carrying the following plasmids: empty vector (pUT18C), wild-type RecB ($pRecB_{WT}$), or $RecB_{F1023A}$ mutant ($pRecB_{F1023A}$), all expressed from pUT18C. Complementation was performed in a *ΔrecA ΔrecB* background. Wild-type RecB served as positive control. No significant difference in DNA content was observed for the $RecB_{F1023A}$ mutant compared to wild type in the absence of UV-induced DNA damage. Comparisons were made using ordinary one-way ANOVA with Šidák correction. Lines represent normalized medians from 2-3 biological replicates; dots indicate individual replicates. ns, non-significant; *$P \leq 0.05$; **$P \leq 0.01$.

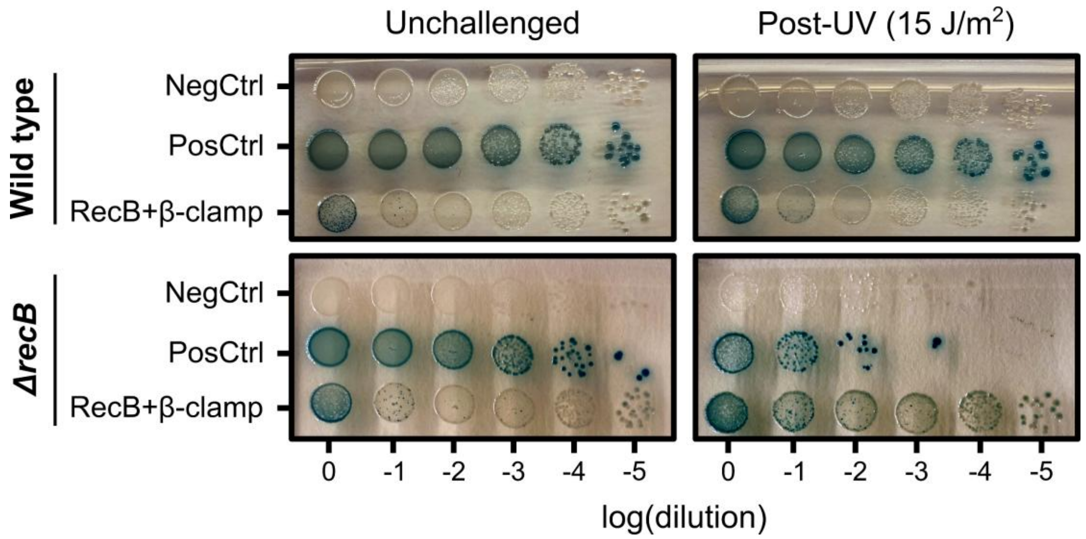


**Figure S5.** UV-induced enhancement of the RecB–β-clamp interaction detected by blue-white assay. Representative blue-white spot assay assessing interaction between RecB–T18 and β-clamp (DnaN)–T25 fusion proteins in wild-type (BTH101, upper panel) and *ΔrecB* (BTH101 *ΔrecB*, lower panel) backgrounds, with (left panels) or without (right panels) UV irradiation (15 J/m²). Empty vector (negative) and leucine zipper (*zip*; positive) controls were included. The assay suggests that the RecB–β-clamp interaction is strengthened following UV-induced DNA damage, particularly in the absence of endogenous RecB.
